# Firewalled synthetic commensal blocks horizontal gene transfer in the gut

**DOI:** 10.64898/2026.09.17.752456

**Authors:** Eric Hu, Kathryn Gorman Durben, Bogdan Budnik, Daniel H. Arlow, Sungwhan F. Oh, Akos Nyerges

**Author notes:** Corresponding author (A.N.).

## Abstract

Synthetic biology enables the rational reprogramming of microorganisms into living therapeutics and agents for bioremediation. However, such genetically modified organisms (GMOs) disseminate their synthetic genetic information into natural microbial communities through horizontal gene transfer (HGT), posing biosafety risks that limit clinical and environmental deployment^1^. Reassigning sense codons to an alternative amino acid identity establishes a genetic firewall that simultaneously prevents incoming and outgoing gene flow^2,3^, but reported implementations compromise fitness, precluding clinical and industrial use. Here, we overcome this limitation using genome design and laboratory evolution to create a high-fitness genetically firewalled *Escherichia coli* commensal. By directly altering the amino acid identity of TCA and TCG serine codons in the genetic code without an unassigned intermediate, we establish a robust genetic firewall that remains stable for thousands of generations. This firewalled commensal stably colonizes the mouse gastrointestinal tract for more than 100 days and blocks viral infections and HGT. As the long-term within-gut evolution of this firewalled organism identified adaptive mutations in genes responsible for carbon source utilization, we rationally redesigned the strain’s genome to increase fitness. Together, this work establishes a genetically firewalled commensal for safer living therapeutics development and provides a strategy for designing high-fitness, virus- and gene-transfer-resistant organisms for clinical and environmental use.

## Introduction

Synthetic biology has transformed our ability to rationally reprogram cells and to use these engineered living organisms as novel therapeutics instead of small-molecule drugs or biologics. Engineered cells harboring recombinant sensors and genetic circuits can control the localization, timing, and dosage of drugs in response to specific disease biomarkers, while protecting their therapeutic cargo from adverse environmental conditions and immune recognition^4^.

However, engineered living organisms inherently proliferate and release their genetic information into natural biomes through horizontal gene transfer (HGT). The recent phase 1/2a clinical trial of Novome Biotechnologies’ NB1000S^5^—an engineered microbial therapeutic that controllably colonizes the human gut—revealed both of these risks: escape from metabolic biocontainment led to persistent gut colonization that could not be cleared by oral antibiotic therapy, while within-patient HGT between the strain and the natural gut microbiome erased the strain’s therapeutic function and spread its engineered colonization module to native gut commensals in at least four of nine patients^6^. Beyond genetic information exchange through conjugation with native microorganisms of the gastrointestinal tract, excreted genetically modified organisms (GMOs) encounter 10^7^–10^10^ bacteriophages per gram of soil and orders-of-magnitude higher numbers in sewage and aquatic ecosystems^1^, which induce viral transduction and facilitate environmental HGT. Prior studies with intentionally released GMOs have highlighted the long-lasting consequences of similar environmental HGT events. In 1989, two engineered *Pseudomonas putida* strains carrying a synthetic phenol-degradation operon were released near a phenol-contaminated Estonian mine^7^. Although the released *P. putida* strains became undetectable within six years, the strains’ synthetic operon spread across indigenous *Pseudomonas* isolates spanning multiple species^7,8^. These case studies indicate that engineered GMOs’ synthetic genetic information can spread readily within native biomes following release.

Consequently, the widespread use of engineered living organisms necessitates the implementation of technologies that prevent the functional escape of synthetic genetic information into the environment. Rational genetic code engineering has emerged as a potential solution to this issue^9–12^. We have previously shown that an artificial genetic code, in which the TCG and TCA (together TCR) serine codons of Syn61Δ3(ev5)^13,14^—an *Escherichia coli* strain in which 98.8% of all annotated instances of genomic TCR and TAG codons were replaced with synonymous alternatives and the corresponding serine tRNA genes (*serU* and *serT*) and RF1 (*prfA*) have been deleted after rounds of random mutagenesis^13,15^—translate as leucine rather than serine due to engineered viral tRNAs, establishes a genetic firewall^3^. This genetic firewall provides isolation against functional incoming and outgoing HGT, including resistance to tRNA-expressing viruses^3,16^. However, existing implementations of this firewall technology drastically reduce fitness, precluding therapeutic and industrial use^2,3,17^. More broadly, despite extensive *in vitro* studies^3,15,18,19^, the behavior of synthetic genomes outside *in vitro* environments remained uninvestigated.

Here, we overcome these limitations to create a high-fitness, broadly virus- and HGT-resistant *E. coli* commensal. By directly swapping the amino acid identity of two serine sense codons in the standard genetic code without an unassigned intermediate and exploiting the deleterious fitness effect of unassigned codons, we addict bacterial cells to artificial translation, establishing an evolutionarily stable, high-fitness genetic firewall. The resulting organism maintains its genetic-code-based firewall *in vitro* for 600 generations and in the gastrointestinal tract of living animals for 102 days, blocks bacteriophage infection *in vitro* and in the mouse gut, and resists gene transfer into natural organisms under laboratory and simulated environmental scenarios. Finally, we redesign our firewalled strain and expand its genome with rationally designed genomic islands from probiotic and commensal *E. coli* strains to confer high-level gut colonization and improve competitive fitness, taking a step toward the clinical translation of genetic code engineered HGT-resistant living therapeutics and the industrial use of virus-resistant organisms.

## Results

### Creation of a high-fitness genetic firewall

We investigated whether directly changing the amino acid identity of sense codons could create a high-fitness genetic firewall. Prior approaches to establish a genetic firewall necessitated the creation of a compressed genetic code by liberating sense codon channels—through the deletion of cognate cellular tRNAs—before reassigning these liberated codon channels to a new amino acid identity^2,3,17^. However, as genome annotation errors and cryptic translated sequences generate unassigned codons on recoded chromosomes, genetic code compression results in a large fitness decrease^9,14,15^. Based on these results, we hypothesized that directly swapping the amino acid identity of sense codons on a recoded chromosome—without first liberating codon channels that could cause ribosome stalling—would maintain fitness and simultaneously addict the recoded organism to its artificial genetic code.

To create a high-fitness firewalled strain and irreversibly addict cells to their modified genetic code, we replaced Syn61 Δ*prfA*’s tRNA^Ser(UGA)^ (encoded by *serT*) with our previously characterized anticodon-swapped bacteriophage-derived tRNA^Leu(CGA)^, which decodes TCG codons as leucine^3^. This tRNA replacement results in cells that partially mistranslate the 92 remaining TCG codons on Syn61’s genome as leucine—as the remaining tRNA^Ser(CGA)^ (*serU*) still decodes TCR codons as serine (**Figure 1a**). Codon identity swap resulted in only an 8.2% fitness decrease compared to the parental strain, indicating that directly changing the amino acid identity of sense codons maintains fitness (**Supplementary Figure 1**).

**Figure 1.**
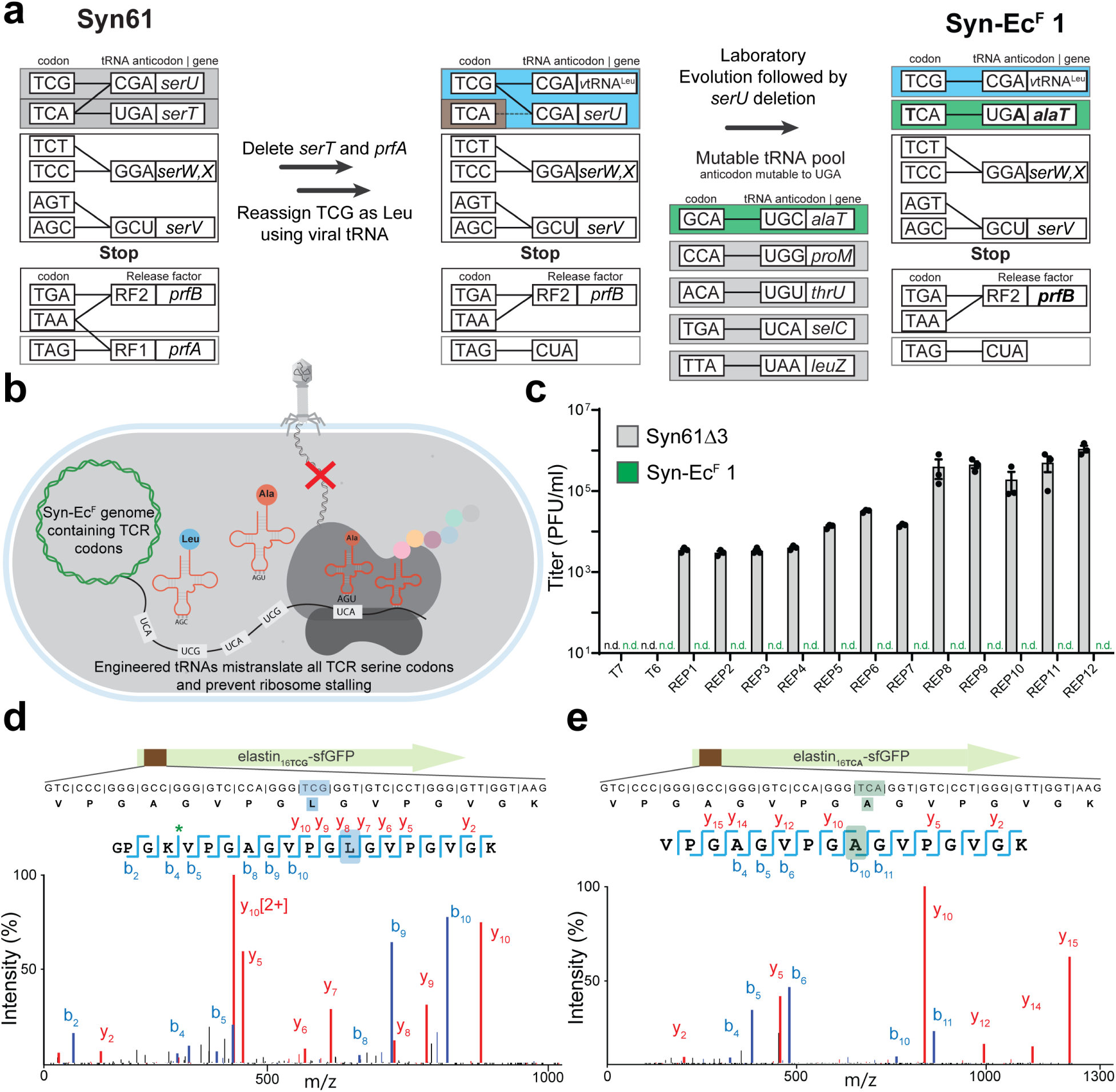
Creation of a high-fitness genetically firewalled organism. (**a**) The creation of Syn-Ec^F^ 1 through the stepwise reassignment of serine TCA and TCG codons. First, we deleted RF1 and replaced Syn61’s tRNA^Ser(UGA)^ encoded by *serT* with an engineered bacteriophage-derived *v*tRNA^Leu(CGA)^. Next, we performed laboratory evolution to adapt cells to their modified genetic code and select an optimal reassignment for TCA codons. Only mutable tRNAs that establish an amino-acid-swapped genetic code are displayed. Finally, we deleted *serU* (tRNA^Ser(CGA)^), eliminating residual serine decoding of TCG codons and creating Syn-Ec^F^ 1. (**b**) Syn-Ec^F^ 1 translates TCG codons as leucine and TCA codons as alanine, and (**c**) confers resistance to an array of laboratory and environmental bacteriophages, including REP1−12, that infect Syn61Δ3^3^. Phage infection assays were performed in *n*=3 independent replicates. Limit of detection = 10 PFU/ml. Bars represent mean bacteriophage titer after 24 hr growth on the corresponding strain, Syn61Δ3 or Syn-Ec^F^ 1; error bars represent standard deviation. (**d**–**e**) The modified genetic code of Syn-Ec^F^ 1 decodes TCG codons as leucine, while TCA codons are translated as alanine. The amino acid identity of the translated TCG codon (**d**) and TCA codon (**e**) within elastin^(16TCG/16TCA)^–sfGFP–His_6_ was confirmed by LC-MS/MS from Syn-Ec^F^ 1 cells. The figure shows the amino acid sequence and LC-MS/MS spectrum of the analyzed elastin^(16TCG/16TCA)^ peptide. Green star indicates a missed trypsin cleavage in the LC-MS/MS-detected peptide.

We next performed Adaptive Laboratory Evolution (ALE), using our established evolution protocol^15^, to adapt cells to their modified genetic code and select an optimal reassignment for the TCA codon channel. As reassigning all TCR serine codons to leucine in Syn61Δ3’s genetic code compromised fitness in our previous work^3^, we relied on ALE to select an optimal amino acid identity for TCA codons from five potential amino acids (**Figure 1a**). As Syn61 contains five tRNAs that are a single mutation away from recognizing TCA codons and can establish an amino-acid-swapped genetic code, and as our ALE workflow explores all single-step mutations across the genome at every ALE step^15^, we hypothesized that tRNA anticodon mutants that can decode TCA codons would readily evolve. In line with this hypothesis, within eight ALE transfers (*i.e.*, 72 generations performed within eight days), laboratory evolution converged on a single variant carrying a mutant tRNA^Ala(UGA)^, induced by an *alaT* anticodon TGC→TGA mutation. This mutant strain translated TCG codons as both leucine and serine, while TCA codons were translated as alanine (**Supplementary Figure 2**). Next, to irreversibly addict cells to their modified genetic code, we deleted *serU* (tRNA^Ser(CGA)^), eliminating the residual serine decoding of TCR codons^3^ (**Figure 1a**).

Finally, to prepare the generated firewalled strain for follow-up *in vivo* experiments, we performed anaerobic gut-mimicking ALE. In this experiment, we performed laboratory evolution in minimal M9 broth supplemented with *E. coli* MDS42’s—and thus, Syn61’s—ancestor, *E. coli* K-12 MG1655’s primary nutrients in the gastrointestinal tract^20^. Following 600 generations performed under gut-mimicking conditions, we isolated a fast-growing variant and subjected it to whole-genome sequencing. Whole-genome sequencing revealed that the evolved firewalled strain accumulated 16 mutations (**Supplementary Table 1**). These included a Thr246→Leu mutation in release factor 2 (RF2), induced by an ACG→TCG codon change, which renders this growth-essential protein dependent on the strain’s altered genetic code and may also alter termination efficiency at the remaining stop codons. We also identified an *rpsG* Leu157→STOP mutation that was shown to restore growth in strains with impaired termination^21^ suggesting that this mutation may compensate for the loss of RF1^22^.

Despite its amino-acid-swapped genetic code that translates TCG codons as leucine and TCA codons as alanine (**Figure 1d,e**), the evolved variant grew faster than the parental Syn61, with a doubling time of 37.2 ±6.3 mins while the parental Syn61 doubles every 50.7 ±6.4 min (*n*=10 independent replicates, mean ± SD, **Supplementary Figure 1**). In comparison, our previous Ec_Syn61Δ3-SL firewalled cells, generated through a codon-compressed intermediate, doubled every 69.3 min^3,16^ while another reported strain with an amino acid-swapped genetic code doubled every 110 min^2^. We named our evolved, high-fitness, firewalled strain Syn-Ec^F^ 1. Next, we confirmed the blocking effect of Syn-Ec^F^ 1’s modified genetic code on bacteriophage replication by repeating our previously described phage replication assay^3^. Bacteriophage resistance tests indicated that Syn-Ec^F^ 1 is resistant to incoming horizontal gene transfer, including resistance to twelve lytic environmental phages known to infect Syn61Δ3 and cellular-tRNA-based amino-acid-swapped-genetic-code-carrying cells by expressing viral tRNA^Ser(UGA)^ (**Figure 1c**)^3^.

Together, these experiments created a horizontal gene transfer-resistant organism stably addicted to its synthetic amino-acid-swapped genetic code and demonstrated that this firewalled strain retains its altered genetic code for at least 600 generations.

### Firewalled cells achieve robust gut colonization

As *Escherichia coli* primarily resides in the lower intestines of warm-blooded animals as a common commensal of the mammalian gut microbiome, we next investigated Syn-Ec^F^ 1’s fitness under gut-mimicking conditions and in the mammalian gut.

We first explored the phenotypic landscape of Syn-Ec^F^ 1, including its ability to occupy metabolic niches important for mammalian gut colonization^20^. We assayed 480 environments, representing the presence of various carbon, nitrogen, sulfur, and phosphorus sources, osmotic and chemical stressors^23^, many of which test essential metabolic functions for intestinal colonization and environmental and gastrointestinal survival^20,24^. Compared to the parental *E. coli* MDS42 strain across all environments, Syn-Ec^F^ 1 exhibited higher fitness, with a cumulative fitness score of 19.5 across all environments. Under identical conditions, Syn61 and Syn61Δ3 displayed a cumulative fitness score of −7.79 and −31.87, respectively, indicating lower fitness under a wide range of conditions. In line with its reduced fitness^15^, Syn61Δ3 failed to grow on minimal M9 agar with glucose as the sole carbon source and lacked the ability to utilize 31% of all carbon sources utilized by the parental MDS42 (**Figure 2a, Supplementary Figure 3**). As *E. coli*’s growth rate in the gastrointestinal tract is carbon source limited^20,24,25^, impaired carbon source utilization suggested a strong negative impact on Syn61Δ3’s *in vivo* growth^20^. In contrast, Syn-Ec^F^ 1 utilized 39% more carbon sources than the parental MDS42 (**Figure 2a,b**) and grew well on minimal M9 agar with glucose as a sole carbon source (**Supplementary Figure 3**).

**Figure 2.**
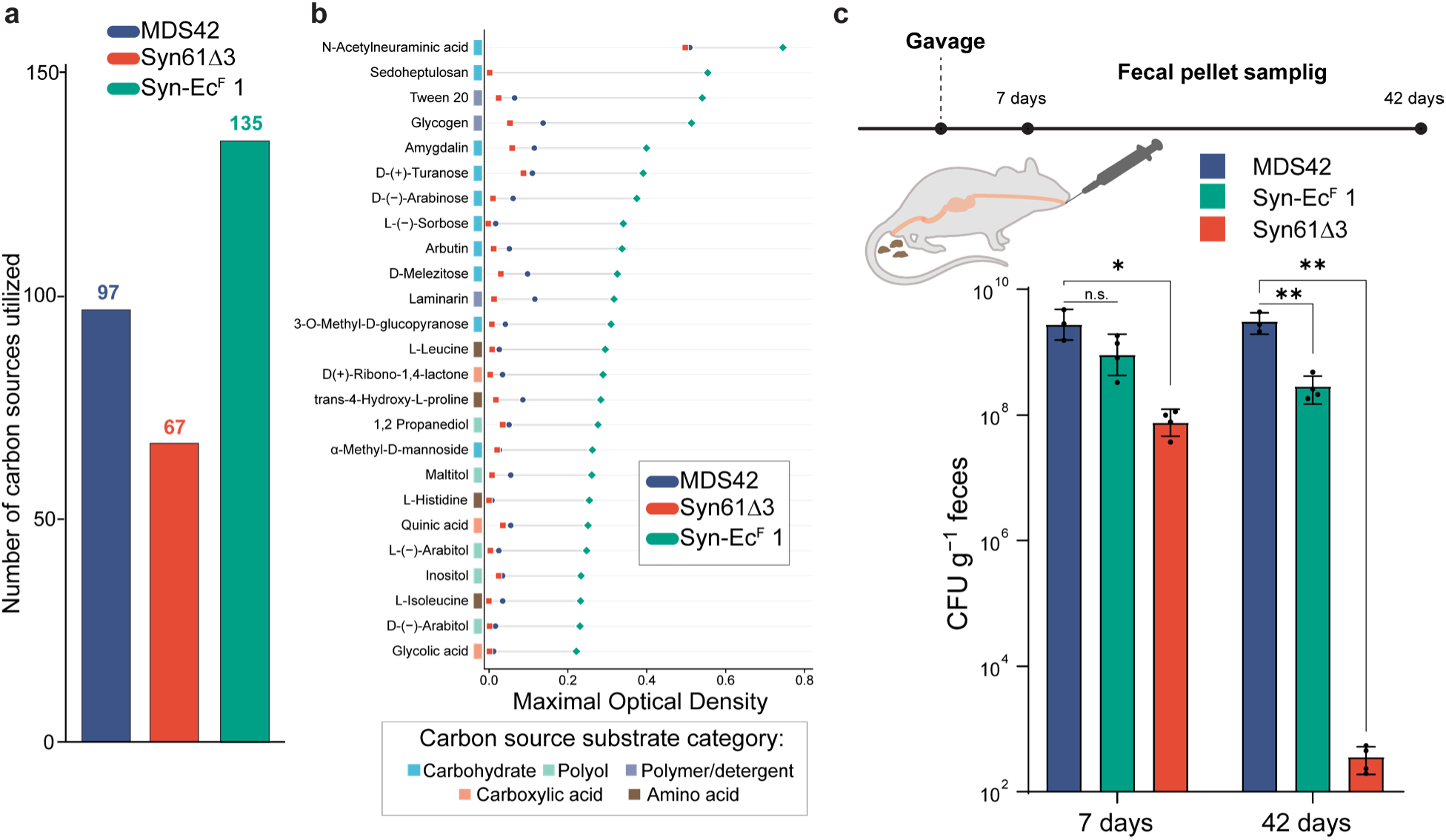
A high-fitness firewalled organism achieves robust gut colonization. (**a**) The number of distinct carbon sources utilized out of 190 conditions tested by *E. coli* MDS42, Syn61Δ3, and Syn-Ec^F^ 1 under identical conditions, measured in two independent replicates using the Biolog Phenotype MicroArray system at 37 °C. Biolog source data are provided in Supplementary Data 2; Biolog data for Syn61Δ3 are described in reference^15^. (**b**) Maximal optical density of *E. coli* MDS42, Syn61Δ3, and Syn-Ec^F^ 1 cultures in the presence of the top 25 carbon sources most efficiently utilized by Syn-Ec^F^ 1 as sole carbon source. (**c**) *In vivo* within-gut fitness of *E. coli* MDS42, Syn61Δ3, and Syn-Ec^F^ 1, 7 days and 42 days after oral gavage of *n*=4 animals. Error bars represent standard deviation; bar graphs represent geometric mean. ** indicates a P < 0.005, * indicates P = 0.0128, while n.s. indicates P > 0.05 based on unpaired two-tailed Student’s t-test.

We next analyzed Syn61Δ3’s and Syn-Ec^F^ 1’s fitness in the mouse gut and compared it to the parental *E. coli* MDS42. To measure *in vivo* fitness in the gut, we gavaged germ-free mice (*n*=4) with 10^7^ colony-forming units (CFU) from each strain separately and housed animals in sterile cages as separate groups. We monitored bacterial gut colonization over six weeks by collecting stool samples and plating homogenates on agar plates. Stool CFU enumeration revealed robust colonization by Syn-Ec^F^ 1 and *E. coli* MDS42, however, Syn-Ec^F^ 1 achieved a lower colonization level than the parental MDS42 (**Figure 2c**). After 7 days in the mouse gut, Syn61Δ3 achieved significantly lower colonization levels than MDS42 and became nearly extinct after six weeks (**Figure 2c**). We hypothesize that Syn61Δ3’s reduced gut colonization is due to a combination of its general fitness defects and its reduced ability to utilize carbon sources available in the murine GI tract.

Phenotypic profiling and mouse gut colonization experiments jointly indicated that directly swapping the amino acid identity of sense codons maintains high fitness for a genetically firewalled organism, both *in vitro* and *in vivo*.

### Firewalled cells block horizontal gene transfer in the mammalian gut

We next investigated the *in vivo* bacteriophage resistance of Syn-Ec^F^ 1 in the mammalian gut. We focused on HGT by bacteriophages as viral transduction is the major driver of *E. coli*’s within-gut evolution and the primary effector of horizontal gene transfer in the gut^26–28^. As before, we colonized germ-free mice with the firewalled Syn-Ec^F^ 1 by gavaging each animal with 10^7^ CFU (**Figure 3a**). As a wild-type control, we colonized animals with *E. coli* Nissle 1917, a widely used probiotic human gut isolate and common host for microbial living therapeutics generation^4,29–31^. Next, after one week of gut colonization, we introduced a mixture of 2×10^8^ plaque-forming units (PFU) of eight lytic bacteriophages via oral gavage into each animal. We simultaneously assayed the effect of lytic and transducing laboratory and environmental phages by introducing the T6 coliphage and six of our recently identified <u>R</u>ecoded <u>E</u>. coli <u>P</u>hages (REP) known to infect Syn61Δ3^3^—*i.e.*, the REP1 and REP3 river water isolates, the REP5, REP7 soil isolates, and the REP9, REP12 pig fecal and farm-derived isolates—plus P1, a widely used laboratory transducing phage. In our *in vivo* phage resistance tests, we targeted a multiplicity of infection (MOI) of 0.1, delivering approximately one bacteriophage per ten bacterial cells in the mouse gut.

**Figure 3.**
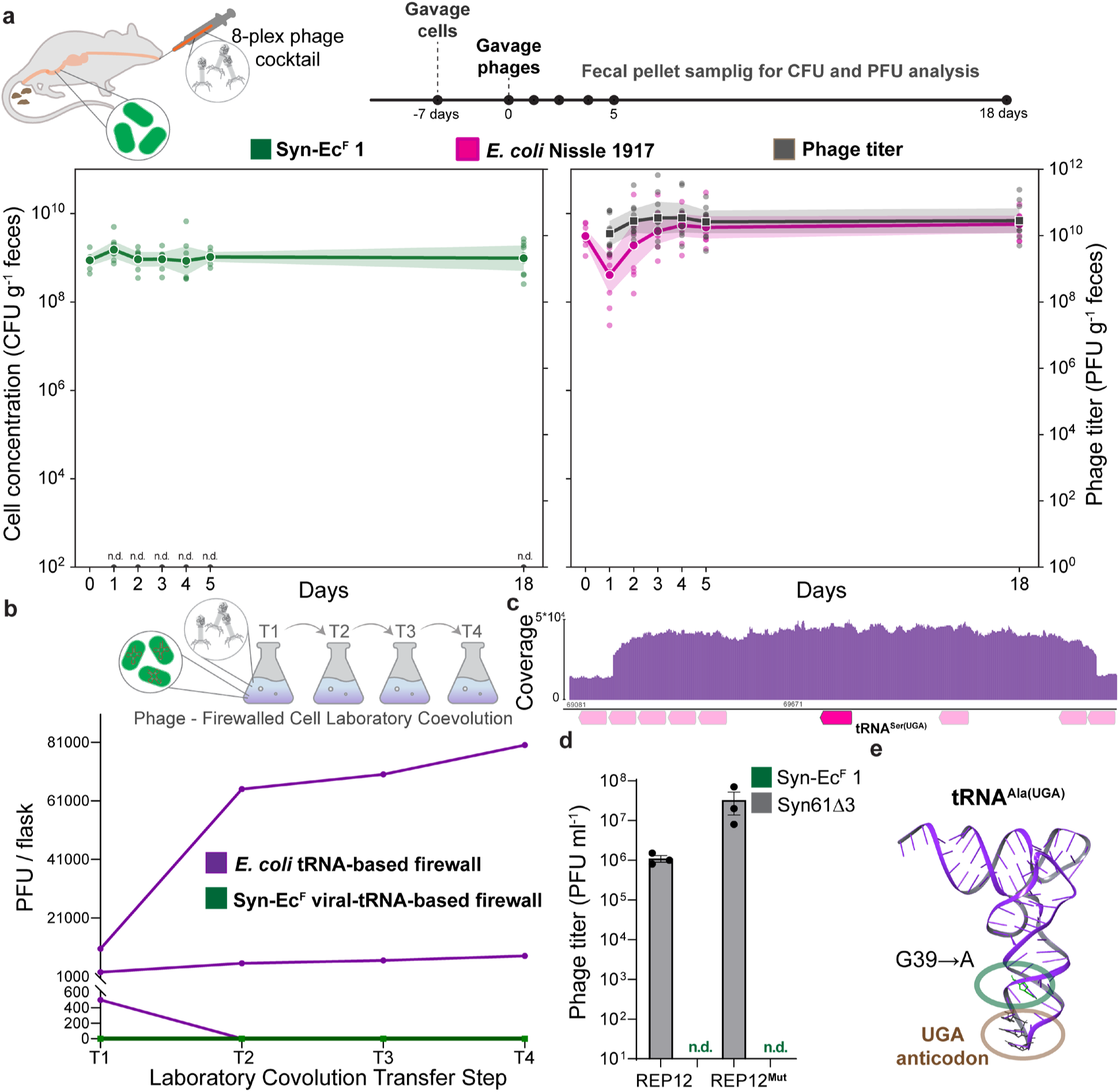
Firewalled organism resists bacteriophages in the murine gut. (**a**) Animals (*n*=10/group) independently pre-colonized with Syn-Ec^F^ 1 and *E. coli* Nissle 1917 were gavaged with an 8-plex mixture of laboratory and environmental bacteriophages, followed by daily fecal sampling for 5 days and then an additional time point at day 18 for bacterial CFU and viral PFU titer evaluation. Plots represent geometric mean; shaded area indicates 95% CI. n.d. represents not detected (limit of detection = 10 PFU/fecal pellet). (**b**) Laboratory co-evolution between bacteriophages and viral-and *E. coli* cellular-tRNA-based amino-acid-swapped genetic code-bearing cells. Liquid cultures of Syn-Ec^F^ 1 and its bacterial-tRNA^Ala(CGA)^-based variant were infected with an 8-plex mixture of laboratory and environmental bacteriophages, followed by transfers and phage titer analysis every 3 days. Phage co-evolution experiments were performed in three independent biological replicates; limit of detection = 10 PFU/ml (*i.e.*, 500 PFU/flask). (**c**) Sequencing read coverage of the REP12 phage mutant (REP12^Mut^) detected in the cellular-tRNA-based firewall + phage co-evolution experiment with a partially triplicated tRNA operon containing the viral tRNA^Ser(UGA)^. tRNAs are marked with magenta; viral tRNA^Ser(UGA)^ is highlighted. (**d**) Bacteriophage titer of the REP12 environmental phage isolate and its evolved derivative (REP12^Mut^) following growth on Syn61Δ3 and Syn-Ec^F^ 1. n.d. indicates not detected; limit of detection = 10 PFU/ml. (**e**) Sequencing the swapped-code-establishing *E. coli* tRNA^Ala(UGA)^ from the co-evolved *E. coli* cells revealed a G39→A anticodon stem-loop mutation within tRNA^Ala(UGA)^. tRNA structure was predicted using AlphaFold 3 and visualized using Schrödinger BioLuminate 2024.3.

Consistent with our prior results and experiments showing that phage titer drops rapidly in co-cultures with firewalled strains (**Figure 1c**)^3^, bacteriophages were undetectable in fecal samples from animals colonized with Syn-Ec^F^ 1 throughout the study (**Figure 3a**) (limit of detection = 10 PFU/fecal pellet), and oral bacteriophage delivery did not affect Syn-Ec^F^ 1 colonization levels.

In contrast, daily time-course phage and bacterial cell counting indicated an immediate drop in *E. coli* Nissle 1917 colonization levels and, simultaneously, high-level phage shedding (**Figure 3a**). A day after phage delivery, animals colonized with *E. coli* Nissle shed lytic bacteriophages at an average of 10^10^ PFU/g feces. In line with prior studies showing that lytic bacteriophages and *E. coli* can stably coexist in the gut microbiome, as *E. coli* primarily replicates in the mucus layer of the gut walls^32^, phage-infected Nissle-colonized animals continued to shed phages throughout the entire colonization study. By the end of Day 5, Nissle-colonized animals’ feces were dominated by members of the *Tequatrovirus* genus, including REP1, REP3, and the T6 coliphage (**Supplementary Figure 4**).

By challenging living animals mono-colonized with a firewalled microorganism, we showed that a genetic-code-based firewall protects against complex mixtures of laboratory and environmental bacteriophages, while a wild-type gut isolate—*E. coli* Nissle 1917—is susceptible to viral predation in the gastrointestinal tract.

### Cellular tRNA-based engineered genetic code does not establish a durable firewall

Following the confirmation of within-gut virus resistance, we explored whether natural viruses could overcome the genetic isolation of firewalled cells over extended periods of time. We have previously shown that establishing an artificial genetic code, in which reprogrammed viral tRNAs translate TCR codons as leucine, creates a genetic firewall that safeguards cells from HGT and infection by tRNA-expressing viruses, but cellular tRNAs cannot establish an efficient firewall^3,16^. Follow-up studies revealed that bacterial phage defense mechanisms often exert their antiviral effect via targeted cellular tRNA cleavage, and bacteriophage-carried tRNAs evolve to avoid recognition by host-defense RNases^33,34^, suggesting that natural evolution may rapidly compromise a cellular-tRNA-based genetic firewall.

To explore the co-evolution of bacteriophages and a firewalled strain bearing either cellular or viral tRNAs, we designed a laboratory viral co-evolution experiment. We initiated adaptive laboratory evolution (ALE) with both Syn-Ec^F^ 1—the viral-tRNA-based firewalled strain— and its derivative in which we replaced the bacteriophage-derived tRNA^Leu(CGA)^ with *E. coli* tRNA^Ala(CGA)^, establishing a previously described bacterial-tRNA-based swapped genetic code^2,17^. In these cells, both TCA and TCG codons are translated as alanine due to the presence of one copy of *E. coli* tRNA^Ala(CGA)^ inserted at the *serT* locus and *E. coli* tRNA^Ala(UGA)^ at the native *alaT* locus.

Next, we infected each evolving ALE population with our previously prepared 8-plex phage mixture at an MOI of 1 and transferred an aliquot of this cell-phage mixture every three days. In line with our prior results demonstrating that cellular-tRNA-based amino acid swapped genetic codes do not establish virus resistance^3,16^, all cellular-tRNA-based cultures showed bacteriophage replication at the start of co-evolution (**Figure 3b**). Within only three ALE transfers, viral titer in the cellular-tRNA-based cultures increased to up to 8×10^4^ PFU/flask. As expected based on published work^3^ and confirmed in our prior tests (**Figure 1c**, **Figure 3a**), viral titer in the viral-tRNA-based Syn-Ec^F^ 1 cultures rapidly dropped below the limit of detection (*i.e.*, 10 PFU/ml) and phages remained undetectable throughout the entire co-evolution experiment (**Figure 3b**).

We isolated individual phages from cellular-tRNA-based cultures and subjected them to Illumina whole-genome sequencing. Genome sequencing revealed the presence of a REP12 phage mutant with a partially triplicated tRNA operon containing the viral tRNA^Ser(UGA)^ (**Figure 3c**). The mutated REP12^Mut^ phage displayed increased titer on Syn61Δ3 but failed to infect the viral-tRNA^Leu(CGA)^-based Syn-Ec^F^ 1 (**Figure 3d**).

Sequencing the swapped-code-establishing *E. coli* tRNA^Ala(YGA)^ from the final time point of co-evolved *E. coli* cells in Flask 2, where viral titers remained low throughout the co-evolution experiment, revealed a G39→A anticodon stem-loop mutation within tRNA^Ala(UGA)^ (**Figure 3e, Supplementary Figure 5**). We note that this tRNA position overlaps with some of the antiphage tRNA nucleases’ and bacterial colicins’ cut site^33–35^. We hypothesize that under strong selection pressure from lytic bacteriophages, *E. coli* tRNA^Ala(UGA)^ evolves to exhibit tRNA nuclease resistance, increasing the intracellular concentration of anticodon-swapped tRNAs and compromising viral translation by competing with phage-produced tRNA^Ser(UGA)^. In contrast to cellular tRNAs, the viral-tRNA^Leu(CGA)^ of Syn-Ec^F^ 1 likely already confers high-level tRNA nuclease resistance, as bacteriophage tRNAs evolve to avoid tRNA nuclease recognition^34^, protecting Syn-Ec^F^ 1 from invading bacteriophages and the effect of tRNA nucleases.

Laboratory co-evolution between a tRNA^Ser(UGA)^-expressing bacteriophage and viral-and cellular-tRNA-based amino-acid-swapped genetic code-bearing cells revealed that a cellular tRNA-based swapped code remains susceptible to bacteriophage evolution while a viral tRNA-based genetic firewall remained stable under these conditions.

### Long-term within-gut evolution of a firewalled organism

We next investigated the long-term stability of Syn-Ec^F^ 1 in the mouse gut, as clinical applications of gut-colonizing microbial living therapeutics necessitate long-term genetic stability^6,36^. Mice are especially suitable for long-term within-gut evolution studies because they recapitulate microbiome evolutionary dynamics and, through coprophagy, exchange microbiota, allowing observation of evolutionary processes with population sizes larger than in individual animals.

As before, we gavaged 19 animals from both sexes with 10^7^ CFU/animal of Syn-Ec^F^ 1 and housed them in a single sterile isolator. Next, we monitored bacterial colonization in each animal’s gut for 102 days post-gavage, approximately 1850 cell generations^37^, by fecal CFU counts every two weeks (**Figure 4a**). CFU measurements indicated that gut colonization levels remained stable throughout the entire colonization study. Based on prior colonization experiments (**Figure 2**) and the average mass of mouse cecal content^38^, we estimated that our mouse colony harbored approximately 10^10^ CFU in total. We confirmed the stability of Syn-Ec^F^’s genetic code and HGT resistance after gut evolution by repeating whole-genome sequencing and our bacteriophage resistance assay on four independent isolates. We observed no anticodon mutations in sequenced isolates. Consistent with no changes in strains’ genetic code, gut-evolved Syn-Ec^F^ retained its high-level phage resistance, and we could not observe phage replication (*i.e.*, phage titers below the limit of detection, *i.e.*, 10 PFU/ml, in supernatant of cultures infected by any of the REP1–12 phages in 3 replicates).

**Figure 4.**
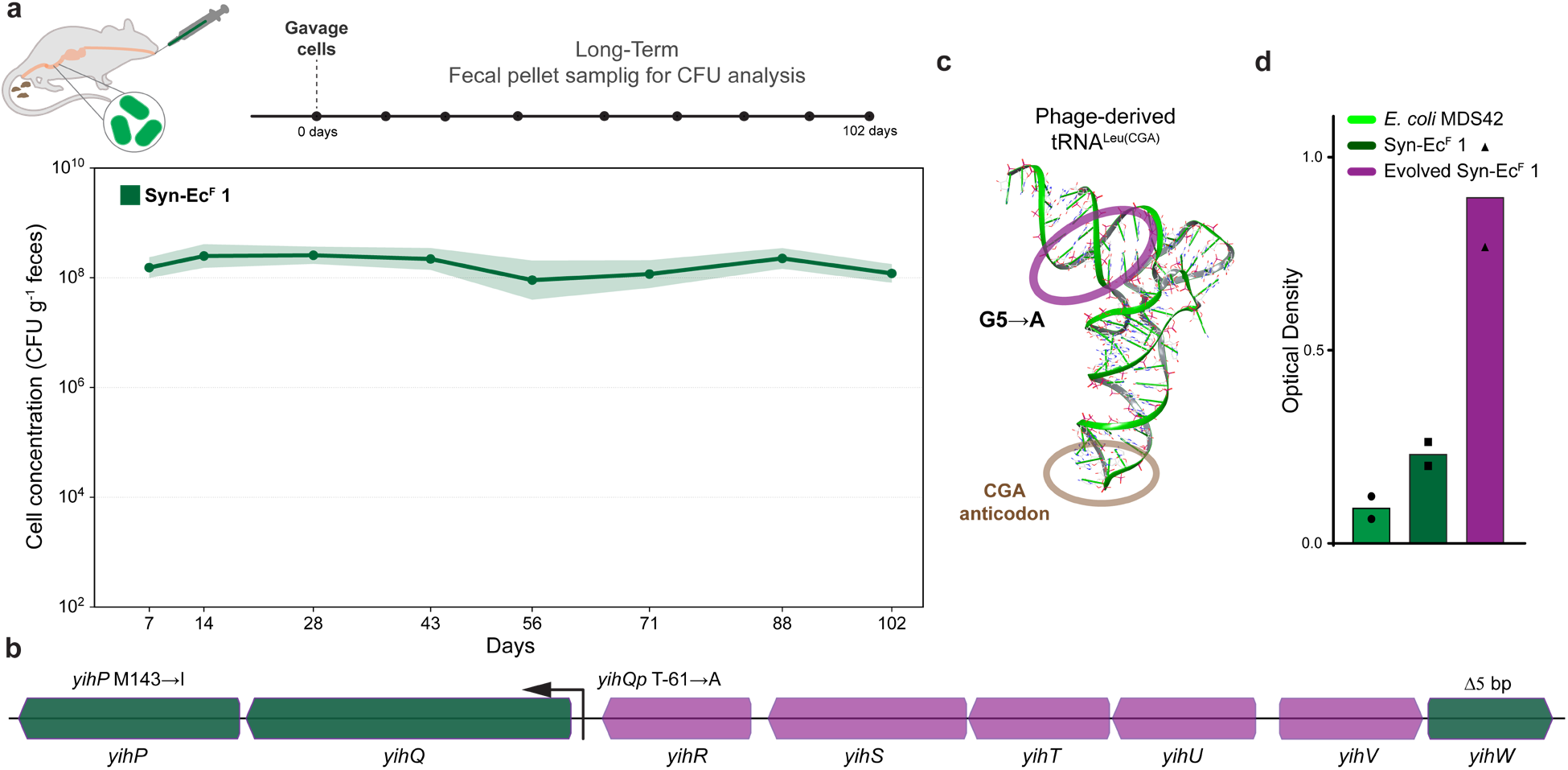
Long-term within-gut evolution of a firewalled organism. (**a**) Long-term stability of Syn-Ec^F^ 1 in the mouse gut. After gavaging *n*=19 animals with 10^7^ CFU/animal of Syn-Ec^F^ 1, fecal bacterial cell counts were monitored for 102 days. Plot represents geometric mean; shaded area indicates 95% CI. (**b**) Genes responsible for sulfoquinovose utilization in *E. coli* K-12 MG1655 and the location of mutations observed in Syn-Ec^F^ 1 following long-term gut evolution. Green arrows indicate mutated ORFs, while magenta arrows mark genes not mutated in the evolved variant. Observed mutations are highlighted; the *yihQ* intergenic promoter mutation is indicated by the position of the *yihQ*p T-61→A and the affected *yihQ*. (**c**) Location of the G5→A acceptor stem mutation in the swapped-code-establishing tRNA^Leu(CGA)^ following the long-term within-gut evolution of Syn-Ec^F^ 1. tRNA structure was predicted using AlphaFold 3 and visualized in Schrödinger BioLuminate 2024.3. (**d**) Growth of *E. coli* MDS42 and the parental and gut-evolved Syn-Ec^F^ 1 on sucrose as the sole carbon source. Bar graph shows the mean of the maximal attained OD_600_, based on *n*=2 independent replicates at 37 °C.

Whole-genome sequencing followed by mutation analysis indicated the acquisition of five mutations across Syn-Ec^F^ 1’s genome, of which four resided in *yihW*, the *yihR*–*yihP* locus, and *dksA,* a major regulator of *E. coli* nutritional response. Three mutations affected CsqR (YihW), the DNA-binding transcriptional regulator of the *yihR*–*yihP* operon, the *yihQ* promoter, and *yihP* (**Figure 4b**). As *E. coli*’s growth rate in the GI tract is primarily limited by carbon-source availability^24,25^ and the *yihR*–*yihP* locus is responsible for sulfoquinovose degradation^39^, an abundant plant-derived sugar and constituent of the mouse feed, we hypothesize that these mutations increase carbon-source availability for Syn-Ec^F^ 1 in the mammalian gut. The remaining mutation occurred in the acceptor stem of the swapped-code-establishing tRNA^Leu(CGA)^ (**Figure 4c**), potentially fine-tuning the interaction of the bacteriophage-derived tRNA with the bacterial translational apparatus.

Finally, we also investigated the effect of accumulated mutations on gut-evolved Syn-Ec^F^ 1’s carbon-source utilization by profiling carbon-source utilization across 190 distinct nutrients, many of which tested metabolic functions for intestinal colonization, environmental and gastrointestinal survival^20,40^. Carbon-source utilization profiling revealed that gut-adapted Syn-Ec^F^ 1 gained the ability to grow on sucrose (**Figure 4d**), a function absent in the parental *E. coli* MG1655^25^, MDS42, or Syn61 strains. As sucrose is one of the most abundant sugars in the mouse feed, we hypothesize that the strong selection pressure caused by the presence of sucrose during intestinal colonization resulted in the *de novo* emergence of sucrose utilization, likely from a low-level promiscuous activity of an enzyme in the sulfoquinovose degradation pathway (**Figure 4b**)^41^. This observation is in line with prior studies showing the convergent selection for sucrose utilization during the within-gut evolution of *E. coli* strains^42,43^.

Long-term adaptive evolution of Syn-Ec^F^ 1 in the mouse gut demonstrated that the viral-tRNA-based genetic firewall is stable and maintains bacteriophage resistance. In combination with our prior *in vitro* evolution, we estimate that Syn-Ec^F^ 1’s genetic code remains conserved for at least 2450 generations. Within-gut adaptation revealed that Syn-Ec^F^ 1 primarily evolves to increase carbon source utilization, suggesting that additional metabolic fine-tuning can further increase its fitness in the mammalian gut.

### Rational genome expansion creates a firewalled commensal

Long-term within-gut evolution indicated selection for mutations in genes responsible for carbon source utilization. Therefore, we rationally redesigned Syn-Ec^F^’s genome to further increase its fitness in the mammalian gut. As the parental Syn61 is based on *E. coli* MDS42, a genome-reduced *E. coli* K-12 MG1655 variant whose genome was rationally minimized to eliminate horizontally transferred genomic regions, including prophages, metabolic functions, and sugar utilization dispensable under laboratory conditions^44,45^, we hypothesized that the elimination of these genomic islands limits Syn-Ec^F^ 1’s *in vivo* fitness.

Based on *E. coli* MDS42’s genome design and the importance of colonization factors necessary for stable gut colonization in closely related *E. coli* commensals (*i.e.*, *E. coli* K-12 MG1655, *E. coli* Nissle 1917, and *E. coli* HS, a nonpathogenic commensal shown to colonize humans at 10^10^ CFU/g of feces)^20,40,44,46,47^, we designed synthetic genomic islands encoding the most important colonization factors absent from MDS42’s genome. These synthetic genomic islands encompassed 1.) metabolic functions essential for *E. coli* K-12 MG1655’s gut colonization^20^, including the ability to import sugars, utilize glycolate and galactitol, 2.) the small regulatory RNA FnrS, deleted during the construction of MDS42, 3.) capsule biosynthesis genes of *E. coli* HS for increased survival in the presence of digestive enzymes, bile, and immune effectors of the GI tract, 4.) high-affinity iron uptake systems via heme utilization and *E. coli* Nissle 1917’s salmochelin siderophore, and 5.) fimbriae biogenesis of *E. coli* HS. Finally, to increase competitive fitness within the gut, 6.) we have also added the probiotic *E. coli* Nissle 1917’s microcin H47 production operon, including its immunity protein. Microcin H47, an antimicrobial peptide produced by *E. coli* commensals, displays *in vivo* inhibitory activity against enteric pathogens, contributing to *E. coli* Nissle 1917’s probiotic effect and increased competitive fitness^48^.

Following our SynOMICS omics-guided genome design workflow^15^, we disentangled overlapping genes and preserved promoter regions while eliminating mobile genetic elements. Next, we recoded the 93 kbp sequence as two 43-and 50-kb segments to match Syn-Ec^F^’s synthetic genetic code by replacing all TCR and TAG codons with synonymous alternatives. We synthesized the DNA as bacterial artificial chromosomes and integrated the synthetic genomic islands into Syn-EcF 1’s genome to produce Syn-Ec^F^ 2 (**Figure 5a**). We confirmed Syn-Ec^F^ 2’s virus resistance by repeating our standard phage replication assay with REP1–12 phages (**Figure 1c**), and we could not observe phage replication (*i.e.*, phage titers below the limit of detection, *i.e.*, 10 PFU/ml, in the supernatant of cultures infected in 3 replicates).

**Figure 5.**
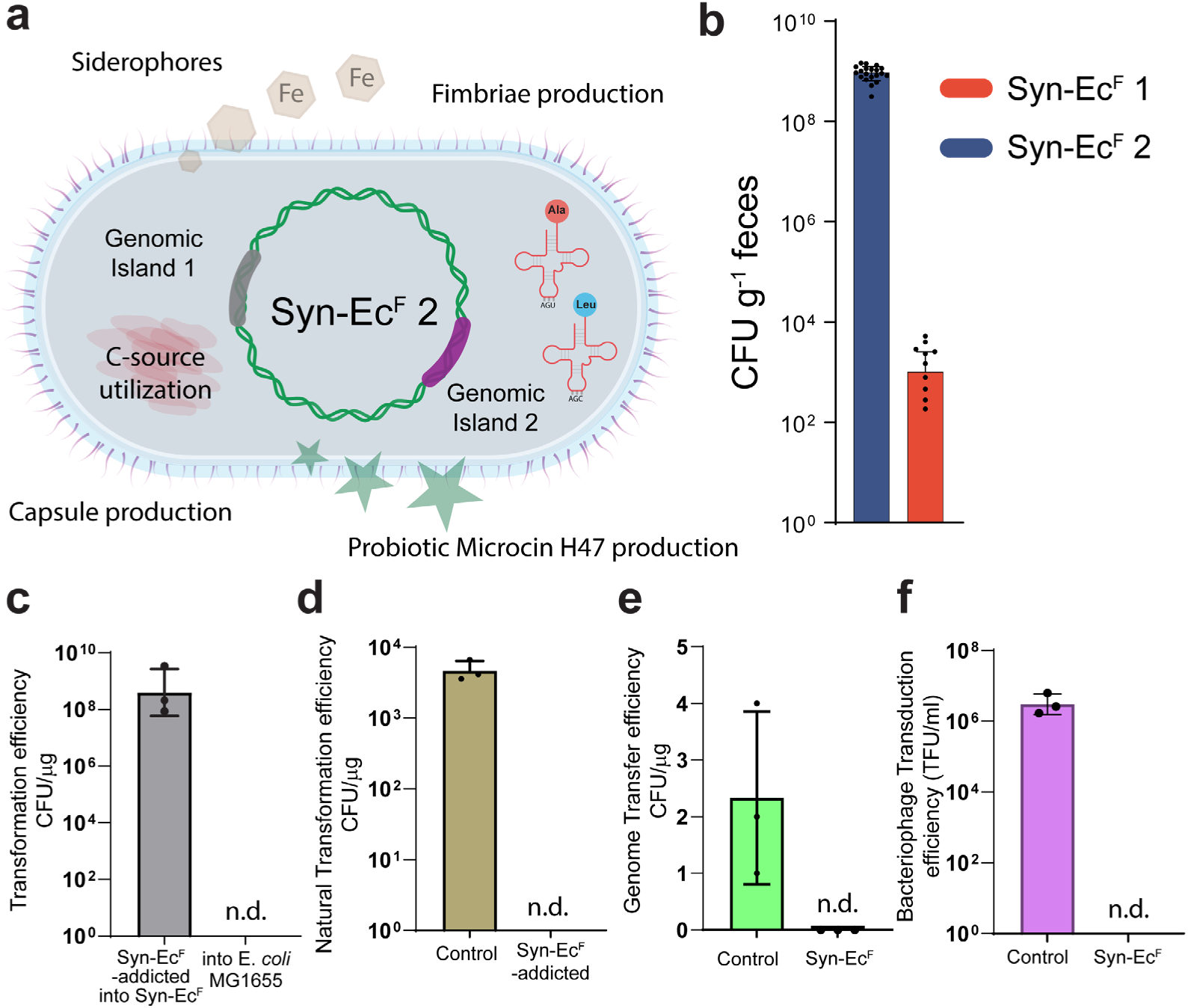
Rational genome design to increase Syn-Ec^F^’s competitive fitness while its engineered genetic code blocks gene transfer. (**a**) We computationally designed, synthesized, and inserted into the genome of Syn-Ec^F^ 1 synthetic genomic islands encoding the most important colonization factors absent from MDS42’s genome. Genomic islands encompassed metabolic functions essential for *E. coli* K-12 MG1655’s gut colonization, the small regulatory RNA FnrS, capsule biosynthesis genes of *E. coli* HS, a gut-colonizing *E. coli* commensal, high-affinity iron uptake systems via heme utilization and *E. coli Nissle* 1917’s salmochelin siderophore, fimbriae biogenesis genes of *E. coli* HS, and the probiotic *E. coli Nissle* 1917’s microcin H47 production operon. (**b**) Increased competitive fitness of Syn-Ec^F^ 2 in the mouse gut. Rational genome expansion of Syn-Ec^F^ 1 yielded Syn-Ec^F^ 2, and we gavaged *n*=20 animals with a 1:1 mixture of both strains. Fecal CFU enumeration after 10 days of gut colonization indicates that Syn-Ec^F^ 2 outcompetes the parental Syn-Ec^F^ 1 strain. Bar graphs represent geometric mean based on *n*=20, error bars represent SD. (**c**) The transformation of the pLS-TCG plasmid into wild-type *E. coli* K-12 MG1655 cells resulted in no escapees. In each case, 3 μg of pLS-TCG plasmid was electroporated into *E. coli* K-12 MG1655 and Syn-Ec^F^ 2 and then plated on antibiotic-containing agar plates. Experiments were performed in triplicate; data points represent data from independent experiments. (**d**) Natural transformation efficiency of the Syn-Ec^F^-addicted pLS-TCG and a control plasmid into *Acinetobacter baylyi* ADP1. (**e**) Genome mass transfer of Syn-Ec^F^ 2-addicted *asd* into *E. coli* Δ*asd* and wild-type *asd* into *E. coli* Δ*asd.* (**f**) T7 phage transduction efficiency into *E. coli* MDS42 (control) and Syn-Ec^F^ 2. Experiments were performed in triplicate; data points represent TFU/ml (transduction efficiency) data from independent experiments.

Next, we evaluated Syn-Ec^F^ 2’s *in vivo* competitive fitness by gavaging 20 animals with a 10^7^ CFU/animal 1:1 mixture of Syn-Ec^F^ 1 and Syn-Ec^F^ 2. After ten days, we collected fecal pellets from all animals and quantified gut colonization levels of both Syn-Ec^F^ strains. In line with its rationally increased competitive fitness, Syn-Ec^F^ 2 rapidly outcompeted Syn-Ec^F^ 1. While Syn-Ec^F^ 2 displayed robust *in vivo* growth with CFU levels reaching up to 1.5×10^9^ CFU/g feces, the parental Syn-Ec^F^ 1’s colonization level dropped to an average of 10^3^ CFU/g feces after ten days (**Figure 5b**).

With rational genome design, we synthetically merged colonization factors and metabolic functions from probiotic and commensal *E. coli* isolates to create Syn-Ec^F^ 2, a synthetic commensal and genomic chimera of three *E. coli* strains that confers high-level gut colonization and shows improved competitive fitness in the gut.

### Firewalled commensal bidirectionally prevents gene transfer

Because the widespread use of engineered living microbial therapeutics requires preventing gene transfer into commensals, pathogens, and members of the native microbiome, we characterized the bidirectional gene transfer frequency of Syn-Ec^F^ 2.

To stress-test gene transfer resistance, we created a Syn-Ec^F^-addicted plasmid vector^3^ (termed pLS-TCG) that depends on 13 TCG—naturally serine-meaning—codons to express leucine in its chloramphenicol-acetyltransferase marker gene, encoded by *cat*. Because of its modified genetic code, pLS-TCG renders Syn-Ec^F^ 1 and 2 cells chloramphenicol-resistant by translating TCG codons as leucine (**Figure 1a**) and expressing functional chloramphenicol-acetyltransferase. At the same time, escape of the same pLS plasmid into cells carrying the canonical genetic code results in the translation of TCG codons as serine and thus the expression of a nonfunctional chloramphenicol-acetyltransferase protein^3^. Next, we mimicked the environmental escape of pLS-TCG by i.) electroporating 3 µg plasmid into 1.3×10^10^ CFU wild-type *E. coli* K-12 MG1655 cells in three replicates followed by plating on chloramphenicol-containing agar plates, and separately, by ii.) incubating 3 µg pLS-TCG plasmid with 6.7×10^9^ CFU of a naturally competent soil isolate, *Acinetobacter baylyi* ADP1^49,50^ in three replicates. Plasmid-mixed *A. baylyi* and electroporated *E. coli* K-12 MG1655 cells yielded no chloramphenicol-resistant escapees carrying the pLS-TCG plasmid (**Figure 5c,d**), whereas the electroporation efficiency of the recipient MG1655 cells was 4.2×10^9^ CFU/µg (*n*=3 independent experiments; based on a pUC-Kan^R^ control plasmid^3^) and *A. baylyi* cells mixed with a control plasmid yielded 4.8×10^3^ CFU/µg (*n*=3 independent experiments). These experiments reinforced our prior results showing that plasmids addicted to an artificial genetic code cannot function in natural cells^2,3^.

We next tested Syn-Ec^F^ 2’s frequency of chromosomal gene escape. This test is especially important given clinical trial results with Novome Biotechnologies’ NB1000S engineered microbial therapeutic^6^, where within-patient bidirectional HGT spread the strain’s engineered genetic information into members of the native human microbiome, causing unpredictable long-term effects and potential environmental dissemination^6,7^. To assay Syn-Ec^F^ 2’s chromosomal gene transfer frequency, we measured the transfer frequency of Syn-Ec^F^’s aspartate-semialdehyde dehydrogenase gene, encoded by *asd—*a frequently used biocontainment gene target for living therapeutics whose deletion renders cells auxotrophic for diaminopimelic acid. We first replaced 9 leucine codons of Syn-Ec^F^ 2’s genomic aspartate-semialdehyde dehydrogenase gene with TCG codons. As Syn-Ec^F^ 2 efficiently translates TCG serine codons as leucine, engineered cells did not require diaminopimelic acid to survive. Next, we simulated environmental gene escape by performing genome mass transfer into recombinogenic recipient cells^51^. The electroporation of 10 µg Syn-Ec^F^ 2 genomic DNA corresponding to 2.3×10^9^ genomes into 10^9^ recombinogenic *E. coli* Δ*asd* recipient cells bearing the standard genetic code, followed by growth-based selection in the absence of diaminopimelic acid, yielded no *asd+* escapees carrying Syn-Ec^F^’s *asd* variant. As a control, we electroporated 10 µg *E. coli* MDS42 genomic DNA, yielding on average 2.3 recombinants/µg genomic DNA that restored *asd* and lost diaminopimelic acid dependency (**Figure 5e**).

Beyond the probability of genomic information exchange with native organisms through direct genome transfer, following excretion, up to 10^10^ bacteriophages per gram of soil and orders-of-magnitude higher numbers in sewage attack GMOs^1^, causing viral transduction and facilitating HGT. Therefore, finally, we have also evaluated the probability of bacteriophage transduction of an antibiotic resistance determinant into Syn-Ec^F^ 2. Because the genomes of Syn-Ec^F^ 1 and 2 both lack genomic prophages and mobile genetic elements, and both strains are resistant to environmental bacteriophages (**Figure 1b,c**), we focused on incoming gene transfer. To measure incoming bacteriophage transduction, we created transducing T7 phages carrying an *E. coli* origin-of-replication and the *tetA* tetracycline resistance gene—containing 16 TCR codons—from transposon Tn10, a widely used antibiotic resistance marker and engineered transposon. Next, we infected 10^11^ Syn-Ec^F^ 2 cells (*i.e.*, 50 ml culture, *n*=3) with transducing T7 phages at an approximate MOI of 10 and plated cells on antibiotic-containing agar plates. We observed no resistant colonies, indicating that Syn-Ec^F^ 2’s engineered genetic code blocks incoming bacteriophage transduction. In parallel, transduction into the parental MDS42 cells yielded 3.53×10^6^ TFU/ml tetracycline-resistant colonies (**Figure 5f**).

The characterization of the bidirectional gene transfer of Syn-Ec^F^ under *in vitro* laboratory and simulated environmental scenarios indicated that cells broadly resist incoming gene transfer. Simultaneously, the synthetic addiction of engineered genetic information to Syn-Ec^F^’s artificial genetic code—through encoding leucine as naturally serine-meaning TCG codons—blocks functional genetic information escape.

## Discussion

Previous works established that rationally engineering the standard genetic code can prevent genetic information flow into and out of genetically modified organisms (GMOs), but prior solutions severely impacted fitness, precluding practical use^2,3,14,17^. Here we show that directly reassigning serine sense codons to translate as an alternate amino acid in an organism’s genetic code creates a high-fitness genetic-code-based firewall that blocks HGT, including bacteriophage infection. In contrast to prior approaches that necessitated the creation of intermediate, compressed genetic codes (*i.e.*, genetic codes with fewer than 64 assigned codons) before establishing genetic isolation^2,3,14,17^, our experiments revealed that directly altering the amino acid identity of sense codons without an unassigned intermediate maintains fitness. We then create a fast-growing firewalled organism—Syn-Ec^F^—that maintains high fitness both *in vitro* and in the mammalian gut. Finally, we demonstrate that Syn-Ec^F^’s engineered genetic code remains stable under laboratory conditions and for more than 100 days in the mammalian gut, and rationally merge genomes of three nonpathogenic gut commensals to increase Syn-Ec^F^’s competitive fitness in the gastrointestinal tract.

Beyond HGT with environmental biomes, over the past four decades, dozens of viral contamination cases were documented in industry^52–54^. These events cost US$ millions and can endanger human lives^53^. A few examples include the simian virus contamination in polio vaccines between 1955–1963, bacteriophage contaminations in the early scaleup of the *E. coli* fermentation process to produce human insulin, bacteriophage contamination in 1,3-propanediol production, and the large-scale *Vesivirus* contamination of Genzyme’s plant leading to temporary shutdown and manufacturing crisis with US$100–300 million in lost sales^53–56^. As multiple prokaryotic and eukaryotic genome recoding projects are ongoing with the goal of creating virus-resistant organisms^57–65^, our results may provide a widely applicable strategy toward the creation and industrial and biomedical use of virus-and HGT-resistant variants of these strains.

Beyond safeguarding genomes from unintentional HGT and viruses, our experiments indicate that rationally expanding existing synthetic, recoded genomes with horizontally transferable genomic regions from phylogenetically related strains can achieve functions not available in existing synthetic strains. As members of the *Enterobacterales* order frequently exchange genetic information through horizontal gene transfer, creating an enterobacterial pangenome^45,46^, we expect that the synthetic expansion of existing recoded genomes (*i.e.*, the reduced-genome *E. coli* MDS42-based^45^ Syn61^13^, Syn57^66^, and Ec_Syn57^15^) with functions from the enterobacterial pangenome can enable a wide range of useful applications, including the creation of firewalled organisms for tumor immunotherapy, nitrogen fixation, and plant growth promotion^29,30,67–69^. In follow-up studies, we will engineer reliance on serine codons to produce leucine-requiring proteins in pangenome-derived genomic regions, preventing HGT and exchange with natural microbiome members. To safeguard these high-fitness synthetic organisms and genomic chimeras from unintended escape, as before^3^, we will establish simultaneous dependence on an amino acid not found in nature.

The United States Environmental Protection Agency (EPA) classifies recombinant genes as poorly mobilizable when their horizontal gene transfer frequency is less than 10^-8^ HGT events per recipient^1,70^, a threshold that broadly applies to all environmentally released engineered microorganisms. No functional gene-escape event was detected in any of our assays, placing the functional HGT frequency of engineered genes in which TCG codons translate as leucine at least an order of magnitude below EPA’s limit. In the unlikely event that functional horizontal gene transfer occurs, the source of engineered genetic information becomes easily identifiable^71^ based on the unique codon composition of Syn-Ec^F^-addicted genetic information. Consequently, these synthetic genes would become selectively targetable using sequence-specific gene drives and RNA-guided DNA nucleases^72,73^. Future work will explore the survival of Syn-Ec^F^ in environmental samples and the long-term fate of swapped-code-addicted mobile genetic elements within microbial communities in the human microbiome, sewage, and soil.

Limitations of this study include the use of germ-free mice mono-colonized with a single *E. coli* strain in most experiments. Mono-colonization enabled us to analyze colonization, gene-transfer susceptibility, and within-gut evolution of cells with engineered genetic codes without microbial community effects, but it limited our conclusions in at least two ways. First, the within-gut fitness and colonization densities we report for Syn-Ec^F^ 1 and Syn-Ec^F^ 2 were measured in the absence of nutrient competition and bacterial cross-feeding by natural microbiome members. Second, mono-colonized animals contain no native exogenous HGT donors, and therefore we relied on an *in-vitro*-produced and orally delivered mixture of lytic and lysogenic laboratory and environmental bacteriophages to test within-gut HGT. Follow-up work will investigate Syn-Ec^F^ 2’s behavior and long-term evolution in human-microbiota-colonized animals.

Beyond synthetic genome recoding, as alternative genetic codes, including sense codon reassignment, appear widely among phylogenetically distant organisms, our findings may provide support for the potential natural emergence of alternative decoding^74^. In our experiments, the deletion of Syn61’s tRNA^Ser(UGA)^ (*serT*) and the reassignment of 176 TCR codons conferred only a mild fitness defect. A similar decoding scenario has been shown to evolve at rare AGG and AGA arginine codons in *E. coli*’s genetic code due to temperature-sensitive tRNA^Arg(UCU)^ mutations (e.g., *argU10*(Ts))^75^. As Syn-Ec^F^ displays high fitness despite reduced decoding at TCA codons and simultaneous mistranslation at 176 serine codons across its genome, we hypothesize that codon reassignments can evolve naturally through stepwise tRNA mutations. For example, the recently described natural AGG arginine-to-methionine reassignment in baboon and human fecal *Bacilli*^76^ could have evolved through mutations that first weakened decoding at AGG and then mistranslated arginine AGG codons as methionine through tRNA^Met(CAU)^ anticodon mutations. In line with this hypothesis, the cognate tRNA for the AGG codon in these organisms displays methionine tRNA identity^76^. As the total number of AGG codons in *E. coli* MDS42 is 954, we hypothesize that intermediate states can remain viable under strong selection pressure, *e.g.*, lytic phages of the mammalian gut. One promising target strain for similar direct reassignment is

*Clostridium sporogenes* NCIMB 10696 (GenBank CP009225.1), a nonpathogenic anaerobe whose genome contains only 239 CGG arginine and 787 TCG serine codons. Importantly, as *C. sporogenes* NCIMB 10696’s genome lacks a cognate tRNA^Arg(CCG)^ and tRNA^Ser(CGA)^, respectively, its genetic code already relies on wobble interactions to decode CGG and TCG codons. As *C. sporogenes* NCIMB 10696 is a validated host for creating tumor-targeting living therapeutics^77^, we hypothesize that the direct reassignment of CGG and/or TCG codons to translate as an alternative amino acid and establish HGT resistance may be feasible. These results—in combination with accelerated experimental evolution^15,78,79^ and steps described in this work—may provide a strategy for creating firewalled organisms without whole-genome recoding and genome synthesis.

Taken together, the bacteriophage-and horizontal-gene-transfer-resistant organism developed in this work, together with the ability to rapidly tailor existing synthetic genomes through genome expansion, is expected to enable a wide range of useful applications. In future studies, we will utilize Syn-Ec^F^ to generate living therapeutics that secrete peptide and protein drugs with an expanded monomer alphabet and create gene-transfer-resistant biocontained organisms for environmental bioremediation, biosensing, and industrial bioproduction.

## Acknowledgments

Funding for this research was provided by the U.S. National Institutes of Health (NIH) and the National Institute of Biomedical Imaging and Bioengineering (NIBIB) under award number R00EB035165 (to A.N). We thank Michael Baym and James J. Collins for useful discussions, Michael Baym and the Baym lab for sharing *A. baylyi* ADP1, and Scott G. Kennedy for helpful comments on the manuscript. We are thankful to Jessica Kay Lang for help with mouse experiments, Biolog, Inc. for support with phenotypic assays, and members of SeqCenter, LLC, Pittsburgh, PA, for support with DNA sequencing experiments.

## Conflict of interest statement

The authors declare competing financial interests. Harvard Medical School has filed provisional patent applications related to the amino acid swapped genetic code in this work, on which A.N. is listed as an inventor. D.H.A. is employed by Ansa Biotechnologies, Inc., but the company had no role in designing or executing experiments.

## Author contributions

A.N. conceived and designed experiments and performed most assays. E.H. performed computational genome design. A.N., S.F.O. and K.G.D. performed mouse assays. B.B. processed proteomics data. D.H.A. coordinated and performed DNA synthesis. A.N. analyzed data and wrote the manuscript with input from all authors.

## Corresponding author

Correspondence to Akos Nyerges.

## Data availability

All strains, sequences, and synthetic DNA constructs utilized in this study are listed in **Supplementary Data 1**. Biolog Phenotypic MicroArray source data is available in **Supplementary Data 2**. Raw data from whole-genome and gut metagenome sequencing experiments will be deposited to the Sequence Read Archive before the publication of the final peer-reviewed work. Further inquiries and material requests should be directed to the lead contact, Akos Nyerges.

## Methods

### Bacterial media and reagents

Gut Mimicking Adaptive Laboratory Evolution experiments used minimal M9 broth supplemented with 0.4% concentration of an equimolar mixture of filter-sterilized sodium gluconate, mannose, N-Acetylglucosamine (GlcNAc), arabinose, ribose, and galactose. All bacterial cell culture and genome editing experiments used 2×YT media consisting of 16 g/l casein digest peptone, 10 g/l yeast extract, and 5 g/l sodium chloride. 2×YT agar plates were prepared by supplementing 2×YT with agar at 1.6% w/v before autoclaving. Lysogeny Broth Lennox (LBL) was prepared by dissolving 10 g/l tryptone, 5 g/l yeast extract, and 5 g/l sodium chloride in deionized H2O and sterilized by autoclaving. Minimal M9 broth with 2% glucose and 0.5 µg/ml thiamine was purchased from Teknova (M8000). Super Optimal Broth (SOB) was prepared by dissolving 20 g/l tryptone, 5 g/l yeast extract, 0.5 g/l sodium chloride, 2.4 g/l magnesium sulfate, and 0.186 g/l potassium chloride in deionized H_2_O and sterilized by autoclaving. Terrific Broth was prepared by dissolving DIFCO™ Terrific Broth (SKU: 243820, Becton, Dickinson and Company) according to the manufacturer’s instructions in deionized H_2_O and sterilized by autoclaving. Top agar for agar overlay assays was prepared by supplementing LB Lennox (LBL) broth with agarose at 0.7% w/v before autoclaving. SM Buffer, 50 mM Tris-HCl (pH 7.5), 100 mM NaCl, 8 mM MgSO_4_, 0.01% gelatin, was used for storing and diluting bacteriophage stocks (Geno Technology, Inc., St. Louis, MO, USA).

### Mice and fecal phage and bacterial cell count analysis

All animal procedures were supervised by the Harvard University Gnotobiotic Core Facility and approved by the Harvard Medical Area Standing Committee on Animals under IACUC protocol IS00003403-3. Experimental animal groups were age and sex matched. Experiments utilized 6-to 8-weeks-old germ-free C57BL/6 mice and all experiments utilized both male and female mice. Mice were housed under 12-hour light-dark cycle and controlled climate (temperature: 21 °C, humidity: 50%) in inflatable plastic positive-pressure sterile isolators. Germ-free C57BL/6 mice were bred and maintained in inflatable plastic isolators on LabDiet 5010 Laboratory Autoclavable Rodent Diet, sterilized by autoclaving, provided *ad libitum*. To facilitate gut microbiome exchange between individually housed groups within the same isolator, during long-term within-gut evolution experiments, we periodically mixed and redistributed animal bedding containing fecal pellets within the same isolator. Bacterial cells were delivered by oral gavage in 100 µl suspension, prepared from early stationary phase cultures grown at 37 °C in 2×YT broth. For bacteriophage administration, mice received bacteriophage mixture in 100 µL of 0.1M sodium bicarbonate *per os* to neutralize gastric acid. Bacteriophages were combined immediately prior to administration from freshly titrated stock solutions stored at 4 °C. We focused on the probiotic *E. coli* Nissle 1917 (EcN, an *E. coli* clade B2 commensal) as control for *in vivo* within-gut bacteriophage assays instead of an isogenic control for Syn-Ec^F^ 1 (*i.e.*, the reduced-genome Syn61 with no clinical relevance), because EcN is a non-laboratory-adapted human gut isolate, has a long history of use in humans, and is widely used as a chassis for engineered living microbial therapeutics, including the development of microbial drugs in clinical trials^4,36^. During colonization studies, fecal pellets were collected under sterile conditions into sterile Eppendorf tubes, weighed, and resuspended in Minimal M9 broth with 2% glucose before performing serial dilutions. To determine bacterial cell concentration, fecal homogenates were serially diluted and 100 µl from each sample was plated to 2×YT agar plates followed by incubation aerobically at 37 °C until colony formation. To determine bacteriophage titer, fecal homogenates prepared in M9 broth were serially diluted in SM buffer and 1.5 µl of the diluted fecal samples were applied to 2×YT agar plates, dried and covered with 0.7% top agar seeded with stationary phase culture of MDS42 and 10 mM CaCl_2_ and MgCl_2_. In the case of fecal pellets from animals colonized with Syn-Ec^F^ 1, 100 µl fecal homogenates were mixed directly with MDS42 cells and 0.7% top agar containing 10 mM CaCl_2_ and MgCl_2_ and poured on prewarmed 2×YT agar plates to decrease PFU limit-of-detection to 10 PFU/fecal pellet. Following 18 hours of incubation at 37 °C, plaques were counted to estimate phage concentration.

### Bacteriophage culturing

Bacteriophage stocks were prepared using the modified liquid lysate Phages on Tap protocol in SOB medium^80^. Phage lysates were prepared from single plaques by picking well-isolated phage plaques into SM buffer and then seeding 3−50 ml early exponential phase cultures of *E. coli* MDS42 cells with the resulted phage suspension in SOB supplemented with 10 mM CaCl_2_ and MgCl_2_. Phage infected samples were grown at 37 °C 100 rpm until complete lysis and then sterilized by filtration. Phage lysates were stored at 4 °C in the dark.

### Phage replication assay

Exponential phase cultures (OD_600_ = 0.3) of Syn61Δ3(ev5) (Addgene #174514) and select strains were grown in SOB supplemented with 10 mM CaCl_2_ and MgCl_2_ at 37 °C. Cultures were infected with phage at an MOI of approximately 0.001. Infected cultures were grown at 37 °C with shaking at 100 rpm in a rotor drum. After 18−24 hours, cultures were spun down at 12,000×g for 2 mins and the supernatant was serially diluted in SM buffer to enumerate output phage concentration.

1.5 µl of the diluted samples were applied to 2×YT agar plates, dried and covered with 0.7% top agar seeded with MDS42 cells and 10 mM CaCl_2_ and MgCl_2_. Following 18 hours of incubation at 37 °C, plaques were counted to estimate phage concentration. In the case of firewalled cultures and cultures where no bacteriophage was observed in 1.5 µl undiluted sample, 100 µl culture supernatant was mixed directly with MDS42 cells and 0.7% top agar containing 10 mM CaCl_2_ and MgCl_2_ and poured on prewarmed 2×YT agar plates to decrease PFU limit-of-detection to 10 PFU/ml culture supernatant.

### Fecal total DNA sequencing

We sequenced the total DNA content of *E. coli* Nissle 1917 and bacteriophage colonized animals by extracting total DNA using the Quick-DNA™ Fecal/Soil Microbe Miniprep Kit (Zymo Research Corporation) according to the manufacturer’s instruction and sequenced extracted total DNA at SeqCenter (Pittsburgh, PA, USA). Sequencing libraries were prepared using the Illumina DNA Prep kit and IDT 10 bp UDI indices with a target insert size of 280 bp and sequenced on an Illumina NovaSeq X Plus, producing 150 bp paired-end reads. Demultiplexing, quality control, and adapter trimming were performed with bcl-convert v4.2.4. Reads were then trimmed to Q28 using BBDuk from BBTools and aligned to genomes of *E. coli* Nissle 1917 and all members of the 8-plex infecting bacteriophage mixture (T6 coliphage, our previously identified environmental Recoded E. coli Phages (REP1, REP3, REP5, REP7, REP9, REP12)^3^ and P1 using Bowtie2 2.3.0^81^ in --sensitive-local mode.

### Bacterial genome sequencing and annotation

Genomic DNA from overnight saturated cultures of isogenic bacterial clones was prepared using the MasterPure™ Complete DNA and RNA Purification Kit (Lucigen) according to the manufacturer’s guidelines and sequenced at SeqCenter (Pittsburgh, PA, USA). Sequencing libraries were prepared using the Illumina DNA Prep kit and IDT 10 bp UDI indices with a target insert size of 280 bp and sequenced on an Illumina NovaSeq X Plus, producing 150 bp paired-end reads. Demultiplexing, quality control, and adapter trimming were performed with bcl-convert (v4.2.4). Reads were then trimmed to Q28 using BBDuk from BBTools and aligned to their corresponding reference by using Bowtie2 2.3.0^81^ in --sensitive-local mode. Single-nucleotide polymorphisms (SNPs) and indels were called using breseq (version 0.36.1)^82^. Only variants with a prevalence higher than 75% were voted as mutations. Following variant calling, mutations were also manually inspected within the aligned sequencing reads using Geneious Prime® 2026.1.

### Sequencing-based gut bacteriophage abundance analysis

We quantified the abundance of the members of our 8-plex bacteriophage inoculum using an Illumina NGS-based approach. In brief, we relied on phage-genome-specific k-mers to identify all eight phages in fecal total DNA samples, which included E. coliphage T6 (GenBank NC_054907.1), P1 (GenBank NC_005856.1), REP1, REP3, REP5, REP7, REP9, and REP12^3^.

The use of a k-mer-based workflow was necessary as most of these phages share high sequence homology with each other and the use of phage-specific read mapping allowed us to distinguish closely related phages in the same sample. As REP5 and REP7, and REP9 and REP12 share more than 90% of their 31-mers, we distinguished these pairs using the presence/absence of 419 and 184 sequence substitutions, respectively, identified with nucmer, delta-filter, and show-snps from MUMmer 4.0.1. To achieve specific detection, for each phage genome, we selected sequence-specific 31mers as unique markers. We retained only 31-mers that occurred exactly once in the target phage genome and did not occur in either the *de novo* sequenced genome of the gavaged *E. coli* Nissle 1917, the mouse genome (GenBank GCF_000001635.27), or any of the other seven phages. Filtering resulted in a final k-mer set that contained 2,358-91,799 markers per phage and covered 4.5-97% of each genome. Following total DNA sequencing as described in section Fecal total DNA sequencing, we first mapped all sequencing reads to the *de novo* assembled genome of *E. coli* Nissle 1917 using Bowtie2 2.3.0^81^ in --sensitive-local mode. Next, unmatched paired-end reads were screened against the k-mer marker set with BBDuk from BBTools 40.02 using exact 31-mer matching. A sequencing read was assigned to a phage only if all the following conditions were met: 1.) the read contained at least two markers specific to that phage, and 2.) the read contained no marker from another phage. For all prior analyses, 3.) only bases with Q≥20 quality score were used. Next, assigned reads were mapped to the corresponding phage genome with minimap2 2.31 in short-read mode and the per-bp sequencing depth was calculated with samtools 1.24. A phage was called present when it had at least five specific reads and at least 95% of its marker loci covered across its genome.

### Recoding and construction of synthetic genomic islands

Based on *E. coli* MDS42’s genome design and the importance of colonization factors necessary for stable gut colonization in closely related *E. coli* commensals (*i.e.*, *E. coli* K-12 MG1655, *E. coli* Nissle 1917, and *E. coli* HS), we designed synthetic genomic islands encoding the most important colonization factors absent from MDS42’s genome. We manually selected these synthetic genomic islands from GenBank-deposited genomes of *E. coli* Nissle 1917 (GenBank CP007799.1), *E. coli* HS (GenBank CP000802.1), and the parental *E. coli* K-12 MG1655 (U00096.3) and merged them into synthetic genomic islands. Next, overlapping genes were disentangled while preserving endogenous promoters (based on Cappable-seq data in *E. coli* K-12 MG1655, RNA-seq data in the case of *E. coli* Nissle 1917, and computational predictions in the case of *E. coli* HS) and ribosome binding sites, followed by the deletion of all prophage and mobile-genetic element-derived ORFs, and all protein-coding genes were recoded to match Syn-Ec^F^ 1’s genetic code. Finally, synthetic genomic islands were synthesized by Ansa Biotechnologies, Inc., and delivered as synthetic Bacterial Artificial Chromosomes (BACs) followed by genomic integration as described in section *Genome editing*. We note that as Ansa’s pStable-Ind-Chlor-v1.1 plasmid was not compatible with Syn-Ec^F^ genetic code due to the presence of 1 TAG and 40 TCR codons, we relied on dsDNA recombineering to replace pStable-Ind-Chlor-v1.1’s origin-of-replication and antibiotic resistance marker with a recoded *trfA* gene and corresponding RK2 origin-of-replication and an *aadA* spectinomycin resistance gene before SynOMICS integration.

### Genome editing

To construct Syn-Ec^F^ 1 and engineer the genome of strains with altered genetic codes, relied on our previously described genome editing plasmids that lack TCA, TCG and TAG codons^3,15^. The sequence of genome editing constructs used in this study is available in **Supplementary Data 1**. We deleted RF1 by making cells electrocompetent for Cas9-assisted recombineering and transforming with 2 μl of 100 μM 90-nucleotide-long single-stranded DNA oligonucleotide (5’-TGATATTCCATTATTCCTGCTCGGACAACGCCGCCAGTTGGTTCATTACTCACGCATTTCAGGATCATCGAGCATCATCTGTGCGGTTTC), deleting the ORF of *prfA* while preserving the expression of the downstream *prmC*. Successful edits were selected by co-transforming 1 μg of a variant of the pCRISPR crRNA expression plasmid carrying a *prfA*-targeting guide sequence (5’-TGATCCTGAAATGCGTGAGA) to cleave the unedited genomic *prfA*. Next, the same procedure was repeated to replace the genomic *serT* using dsDNA recombineering with a synthetic cassette encoding the SLP2018-2-101^83^ promoter followed by the bacteriophage-derived (GenBank ID MF402939.1) tRNA^Leu(CGA)^ and a proK terminator, yielding Syn61 Δ*serT* Δ*prfA* MF402939.1 *v*tRNA^Leu(CGA)^. Next, we performed eight ALE transfers with our terascale laboratory evolution protocol^15^ in 2×YT broth and repeated the same ssDNA recombineering procedure to delete *serU*. Genome expansion using SynOMICS utilized Syn-Ec^F^ 1 as base strain and recoded versions of our previously described gentamicin resistance-conferring genomic deletion cassettes^15^ (**Supplementary Data 1**). As genome expansion only expanded the already existing genome of Syn-Ec^F^ without replacing genomic regions, in contrast to the standard SynOMICS process that relies on stepwise deletion of the parental genome, we inserted Synthetic Genomic Island 1 between genes *yihF* and *yihG*, while Synthetic Genomic Island 2 was inserted between genes *yeeJ* and *yeeL* of Syn-Ec^F^ 1. We selected these regions as safe insertion sites based on no mRNA and ribosome coverage in prior multi-omics datasets generated using *E. coli* MDS42 and Syn61Δ3^15^. To integrate synthetic genomic islands, in brief, we co-electroporated 10 μg of a dsDNA cassette carrying a recoded gentamicin-resistance-conferring gene with flanking 400 bp terminal homologies in genes *yihF* and *yihG*, and in genes *yeeJ* and *yeeL* of Syn-Ec^F^ 1, respectively, and 6 μg of a genome-targeting crRNA plasmid that cleaves the unedited genome. Following electroporation, cells were recovered overnight at 32 °C 250 rpm in 2×YT, and 0.5 to 3 ml culture was plated in multiple replicates onto 145×20 mm Petri dishes (Greiner Bio-One) containing 2×YT agar with the corresponding antibiotics and incubated at 32 °C until colony formation. Finally, and as in the standard SynOMICS workflow, colonies were first validated by external and junction-specific PCRs followed by whole-genome sequencing to confirm the desired edits. Next, we repeated induced competent cell preparation followed by the transformation of SynOMICS’s pINTsg plasmid. Following electroporation, cells were recovered overnight at 32 °C 250 rpm in 2×YT containing 1% glucose, and 0.1 to 0.5 ml of culture was plated in multiple replicates onto 145×20 mm Petri dishes (Greiner Bio-One) containing 2×YT agar with the corresponding antibiotics and 1% glucose and incubated at 32 °C until colony formation. Finally, and as in the standard SynOMICS workflow, colonies were first validated by external and junction-specific PCRs that tested the loss of the gentamicin-resistance cassette and the integration of the corresponding genomic island, followed by whole-genome sequencing to confirm the desired edits. We used Syn-Ec^F^ 1, with an integrated gentamicin-resistance cassette that was shown to confer no fitness cost^84^, within the *yihF−yihG* locus, as a competition partner for Syn-Ec^F^ 2.

### Bacteriophage and firewalled cell co-evolution experiments

To explore the co-evolution of a firewalled strain bearing cellular or viral tRNAs and bacteriophages, we initiated laboratory co-evolution with both Syn-Ec^F^ 1 and its derivative in which we replaced the bacteriophage-derived tRNA^Leu(CGA)^ with *E. coli* tRNA^Ala(CGA)^ (5’-GGGGCTATAGCTCAGCTGGGAGAGCGCCTGCTTCGAACGCAGGAGGTCTGCGGTTCGATCCCGCATAGCTCCACCA), establishing a bacterial-tRNA-based swapped genetic code. We utilized dsDNA recombineering to replace Syn-Ec^F^ 1’s bacteriophage-derived tRNA^Leu(CGA)^ with *E. coli* tRNA^Ala(CGA)^ creating an isogenic cellular-tRNA-based version of Syn-Ec^F^ 1 in which TCR serine codons translate as alanine. Next, we infected 50 ml of each evolving early-stationary phase (*i.e.*, OD_600_ = 0.8) ALE population with an equimolar mixture of T6 coliphage, our previously identified environmental Recoded *E. coli* Phages (REP1, REP3, REP5, REP7, REP9, REP12) and P1 at an MOI of 1 and transferred 1 ml aliquot of cell-phage mixture into 50 ml fresh 2×YT every three days. Cultures were incubated at 37 °C, 250 rpm. To determine bacteriophage titer, samples from evolving cultures were pelleted at 14,000×g for 2 mins and serially diluted in SM buffer. 100 µl of the diluted samples were mixed with MDS42 cells and 0.7% top agar containing 10 mM CaCl_2_ and MgCl_2_ and poured onto 2×YT agar plates. Following 18 hr of incubation at 37 °C, plaques were counted to estimate phage concentration. Whole-genome sequencing of the mutated REP12_Mut_ phage was performed by infecting *E. coli* cells with single-plaque isolates prepared as described in section *Bacteriophage culturing* to create a high-titer lysate and extracting total DNA from the culture supernatant with the Quick-DNA™ Fecal/Soil Microbe Miniprep Kit (Zymo Research Corporation) according to the manufacturer’s instructions, and sequenced extracted total DNA at SeqCenter (Pittsburgh, PA, USA). Sequencing of the chromosomal tRNA^Ala(UGA)^ and tRNA^Ala(CGA)^ regions was performed by first PCR amplifying the target region followed by amplicon Nanopore sequencing (Plasmidsaurus, USA). We predicted tRNA structures using AlphaFold v3.0.4 with six magnesium ions as counterions and visualized them in Schrödinger BioLuminate 2024.3.

### Construction of pLS-TCG

The whole pLS-TCG plasmid was synthesized by GenScript in a pCC1 backbone followed by the removal of the pCC1 backbone by whole-plasmid PCR amplification and circularization by ligation with T4 DNA ligase (New England Biolabs, USA). Following pLS-TCG plasmid assembly, purified ligated PCR amplicons were electroporated into Syn-Ec^F^ cells and plated onto chloramphenicol-containing 2×YT agar plates after overnight recovery at 32 °C. Plates were incubated until colony formation at 32 °C. Finally, plasmids from antibiotic-resistant clones were subjected to direct-colony whole-plasmid sequencing (Plasmidsaurus, USA) and aligned to pLS-TCG to confirm the sequence using Geneious Prime® 2026.1.2.

### Escape rate analysis of the pLS-TCG plasmid

We analyzed the ability of pLS-TCG plasmids to function outside Syn-Ec^F^ cells by transforming extracted plasmids into *E. coli* K-12 MG1655, and separately, by mixing plasmids with the naturally competent soil isolate, *Acinetobacter baylyi* ADP1. pLS-TCG plasmids were purified from Syn-Ec^F^ 1 using the PureLink^TM^ Fast Low-Endotoxin Midi Plasmid Purification Kit (Thermo Fisher Scientific, USA) according to the manufacturer’s instructions. Next, we electroporated 3 μg plasmid into freshly made electrocompetent cells of *E. coli* K-12 MG1655. Cells were made electrocompetent by diluting an overnight 2×YT culture of MG1655 1:100 into 500 ml 2×YT in a 2000 ml flask and growing cells aerobically at 32 °C with shaking at 250 rpm. At OD_600_ = 0.3, cells were cooled on ice and then pelleted by centrifugation and resuspended in 10% glycerol-in-water. Cells were washed 4-times with 10% glycerol-in-water and then resuspended in 400 μl 20% glycerol-in-water. 3000 ng from each plasmid sample was then mixed with 80 μl electrocompetent cells and electroporated by using standard settings in two 1-mm electroporation cuvettes by using standard electroporator settings (1.8 kV, 200 Ohm, 25 μF). Electroporations were performed in three replicates. Electroporated cells were then resuspended in 1 ml 2×YT, and the culture was allowed to recover overnight at 37 °C with shaking at 250 rpm. Finally, 2 × 500 μl from each recovery culture was plated to 2×YT agar plates 30 μg/ml chloramphenicol in 145×20 mm Petri dishes (Greiner Bio-One). As a control for electroporation-based escape assays, we electroporated pLS-TCG into Syn-Ec^F^ cells and plated serial dilutions onto chloramphenicol-containing agar plates. Plates were incubated at 32 °C for up to seven days and inspected for growth. Electroporation efficiency measurements were performed by electroporating the same plasmid into the parental Syn-Ec^F^ cells under identical conditions. Natural competence-based uptake was measured by growing *Acinetobacter baylyi* ADP1 cells in LBL at 32 °C in 3 ml media overnight, followed by diluting cells 1:100 in 1000 µl fresh antibiotic-free LBL with either 3 μg pLS-TCG plasmid or an isogenic plasmid containing a nonrecoded *cat* resistance marker. The culture was allowed to recover overnight at 32 °C in a rotor drum, followed by serial dilutions and plating on 2×YT agar plates with 30 μg/ml chloramphenicol. In the case of pLS-TCG, all transformed cells were plated onto 145×20 mm Petri dishes containing 2×YT agar plates with 30 μg/ml chloramphenicol and incubated for 7 days.

### Phage transduction assays

We evaluated Syn-Ec^F^’s frequency of incoming bacteriophage transduction by creating T7 transducing phages. Transducing *T7-tetA* plasmids were generated as described earlier^85–87^. In brief, we *de novo* synthesized the T7 coliphage terminal repeat region containing the endogenous T7 RNApol promoter and subcloned it together with the *tetA* locus of transposon Tn10 from our previously described RK2-FCas7 plasmid^15^. Next, we prepared transducing T7 bacteriophages by preparing high-titer T7 phage lysate from *E. coli* MDS42 cells harboring the *T7-tetA* plasmid as described in the section *Bacteriophage culturing*, but with the following modification: 30 mins before T7 phage addition, cells were pelleted 4000×g for 10 mins and resuspended in prewarmed media containing 2 mM CaCl_2_ and MgCl_2_ to remove antibiotics. Following the addition of 100 ul T7 phage stock at an approximate MOI of 1, cultures were allowed to lyse for 120 mins at 37 °C 100 rpm followed by the removal of cell debris by centrifugation and filtering on a 0.45 µm membrane. Transducing phage preparations were stored for a maximum of one week at 4 °C in the dark. We measured phage transduction efficiency of Syn-Ec^F^ 2 by pelleting late-exponential (*i.e.*, OD_600_ = 0.7) Syn-Ec^F^ 2 cells grown in 2 mM CaCl_2_ and MgCl_2_ 2×YT at 37 °C and resuspending cells in 50 ml undiluted T7 phage lysate from *E. coli* MDS42 cells harboring the *T7-tetA* plasmid to achieve an MOI of 10. Following 15 mins of phage adsorption at 37 °C, cells were allowed to recover for 1 hr at 37 °C under gentle agitation. Finally, all cells were pelleted, resuspended in 2×YT, and plated on ten 14.5 cm 2×YT agar plates containing 15 μg/ml tetracycline and incubated for 3 days at 37 °C. We determined transducing phage titers by measuring transduction efficiencies (TFU/ml), determined by serially diluting T7 phage lysate from *E. coli* MDS42 cells harboring the *T7-tetA* plasmid in SM buffer and mixing diluted samples with equal volumes of late-exponential *E. coli* MDS42 cells, as described before^86^. Following 15 mins of phage absorption at 37 °C, cells were allowed to recover for 1 hr at 37 °C and then plated onto 2×YT agar plates containing 15 μg/ml tetracycline and incubated at 37 °C until colony formation.

### Genome mass transfer-based gene transfer assays

We performed genome mass transfer by transforming freshly extracted (*i.e.*, non-fragmented) gDNA into recombinogenic target cells. Genomic DNA was extracted from stationary-phase cultures using the MasterPure™ Complete DNA and RNA Purification Kit (Lucigen) according to the manufacturer’s guidelines and resuspended in H_2_O. Next, we electroporated 10 µg gDNA (in 3-4 µl) into freshly prepared induced electrocompetent cells of *E. coli* Δ*asd* cells containing the pORTMAGE-2 plasmid that expresses λRed and a dominant negative MutL E32→K allele all controlled by temperature sensitive cI857 repressor (Addgene plasmid #72677), inactivating methyl-directed mismatch repair (MMR) and supplying Lambda phage’s exo, beta, gam. Induced pORTMAGE-expressing cells were prepared and transformed according to our standard protocol^88^ in the presence of 100 µg/ml 2,6-diaminopimelic acid (DAP). Electroporated cells were allowed to recover overnight at 32 °C in the presence of DAP, washed by pelleting cells at 4000xg and resuspending them in DAP-free medium, followed by plating on 2×YT agar plates without DAP. Plates were incubated at 32 °C until colony formation, and DAP-insensitive colonies were identified by restreaking colonies onto fresh DAP-free 2×YT agar plates. Finally, we identified recombinants that reverted the genomic *asd* locus by colony PCR. We note that PCR-based validation was necessary, as the genomic Δ*asd* cells exhibit an escape frequency around 10^-9^ escapee/cell. In our experiments, compensatory mutants that escaped DAP dependence via genomic compensatory mutations outside the *asd* locus were easily identifiable by colony morphology.

### Gut-mimicking Adaptive Laboratory Evolution

We performed anaerobic gut-mimicking adaptive laboratory evolution experiments in minimal M9 medium supplemented with carbon sources primarily responsible for the gut colonization of E. coli K-12 MG1655 in the mouse gut. At each daily transfer step, 1 ml cells were transferred into 500 ml minimal M9 broth supplemented with 0.4% final concentration of an equimolar mixture of filter-sterilized arabinose, sodium gluconate, mannose, N-acetylglucosamine (GlcNAc), ribose, and galactose and incubated anaerobically for 24 hours at 37 °C, 250 rpm in a 2000 ml baffled Erlenmeyer flask with an air-tight cap. We did not intentionally remove oxygen from the culture medium and headspace and relied on the facultative anaerobic *E. coli* cells’ metabolism to utilize oxygen until anaerobic growth was achieved. With these steps, we aimed to mimic industrial cultivation followed by oral delivery and then anaerobic growth in the gut. Cultures were grown until saturation at every ALE step. Adaptive laboratory evolution experiments were performed in two replicates, starting from the same seed culture of the starting strain. During ALE experiments, cells were plated onto 2×YT and M9 glucose agar plates following every five transfers and incubated at 37 °C aerobically to assess growth rate based on the speed of colony formation. Finally, evolution experiments were terminated by spreading bacterial cells onto 2×YT agar plates, incubated at 37 °C, and an individual, fast-growing colony from each experiment was isolated and subjected to whole-genome sequencing.

### Doubling time and phenotype measurements

Growth parameters were determined as previously described^23^. Briefly, to determine growth parameters under standard laboratory conditions, saturated overnight cultures, initiated from isogenic, single colonies, were diluted 1:100 into 100 µl 2×YT in a 96-well flat-bottom transparent microtiter plate (Corning Inc., USA). Overnight starter cultures were grown in the same media as the downstream conditions. To assess growth kinetics, diluted cultures in ten replicates were incubated aerobically at 37 °C, 800 rpm, and 1-mm orbital shaking in a BioTek Synergy H1 Multimode plate reader (Agilent Technologies, Inc., USA). Optical density at 600 nm (OD_600_) measurements were taken every 9 minutes until the stationary phase was reached. Finally, we calculated the doubling time from background-normalized OD_600_ values using the open-source GrowthRates package, version 4.4^89^. Biolog Phenotype microarray measurements were performed on PM Phenotype MicroArrays™ (Biolog) plates PM01, PM02 (carbon sources) and PM04 (phosphorus and sulfur sources), PM06 (nitrogen sources) and PM09 (osmotic and ionic stressors), as previously described, utilizing the same instruments and reagent stocks^23^. In brief, strains were grown overnight at 37 °C from single colonies and diluted 1:500 into 15 ml Biolog inoculation fluid (IF-0, Biolog). PM plates were inoculated with 100 μl cell suspension per well and incubated at 37 °C in an Odin plate reader (Biolog) for 48 hours with kinetic OD monitoring every 20 minutes, based on the manufacturer’s protocol. PM Phenotype MicroArrays™ growth results for Syn61Δ3 were determined under identical conditions in reference^15^ and we utilized the same dataset in this work. Measurements were performed in two independent replicates, baseline-normalized based on the control well on each plate, and OD values were compared to *E. coli* MDS42.

### Elastin_16TCR_-sfGFP-6×HIS expression

We assayed the amino acid identity incorporated at TCG and TCA codons by expressing an MSKGPGKVPGAGVPG**X**GVPGVGKGGGT-elastin peptide fused to sfGFP with a terminal 6×His tag (in which **X** denotes the analyzed codon, TCA or TCG)^90^ on a plasmid containing a gentamicin resistance gene and an RK2 origin-of-replication (**Supplementary Data 1**). For LC/MS-MS measurements, we diluted cultures 1:100 from overnight starters into 30 ml gentamicin-containing BD DIFCO™ Terrific Broth (Becton, Dickinson and Company, #243820) in 300 ml shake flasks and cultivated cells for 36 hours at 37 °C, 250 rpm aerobically. We then determined the elastin peptide’s sequence by pelleting and washing cultures with ice-cold PBS (Phosphate Buffered Saline) and resuspending cell pellets in BugBuster Protein Extraction Reagent (MilliporeSigma) based on the manufacturer’s protocol. The lysed cell mixture was spun down twice at 18,000×g for 17 minutes, and the supernatant was mixed in a 1:2 ratio with HIS-Binding/Wash Buffer (G-Biosciences, USA) and 50 µl HisTag Dynabeads (Thermo Fisher Scientific, USA). Following an incubation period of 5 minutes, the beads were separated on a magnetic rack and washed with 300 µl HIS-Binding/Wash Buffer and PBS. After the last PBS wash step, the bead pellets containing the bound elastin-sfGFP-6×HIS protein samples were frozen at -80 °C until LC-MS/MS sample preparation.

### Liquid chromatography-tandem mass spectrometry (LC/MS-MS) analysis of tryptic elastin-sfGFP-6×HIS

Samples from elastin-sfGFP-6×HIS expression experiments were digested directly on HisTag Dynabeads according to the SP3 digest procedure^91^. In brief, samples were washed with 50 mM TEAB (triethylammonium bicarbonate buffer) and then rehydrated with 50 mM TEAB-trypsin solution, followed by an overnight digest. Digested peptides were then separated from HisTag Dynabeads and concentrated by spinning and drying samples at 3.000×g using a SpeedVac concentrator. Samples were then solubilized in 0.1% formic acid-in-water for subsequent analysis by tandem mass spectrometry. LC-MS/MS analysis of digested samples was performed on a Astral Orbitrap Mass Spectrometer equipped with a NEO nano-HPLC (both from Thermo Fisher Scientific, USA). Peptides were separated on a 300 µm inner diameter microcapillary trapping column packed first with 1 cm of C18 Reprosil resin (2 µm, 100 Å, from Thermo Fisher, MA) followed by a 15 cm 75 µm analytical column (IonOptics, Australia). Separation was achieved by applying a gradient from 4% to 30% acetonitrile in 0.1% formic acid over 24 mins at 250 nl/min. Electrospray ionization was performed by applying a voltage of 2 kV using an IonOptics electrode junction at the front of the analytical column. The mass spectrometry survey scan was performed in the Orbitrap in the range of 400−800 m/z at a resolution of 2.4×10^5^, followed by nDIA window width of 2 m/z, AGC (automatic gain control) setting of 7 ms. The raw data were analyzed using PEAKS 12.5 (Bioinformatics Solutions, Canada). Assignment of MS/MS spectra was performed using the Sequest HT algorithm by searching the data against a protein sequence database, including all protein entries from *E. coli* K-12 MG1655, all protein sequences of interest (including the elastin-sfGFP fusion protein), as well as other known contaminants such as human keratins and common lab contaminants. PEAKS searches were performed using a 10-ppm precursor ion tolerance and requiring each peptides N-/C termini to adhere with trypsin protease specificity while allowing up to two missed cleavages. Methionine oxidation (+15.99492 Da), deamidation (+0.98402 Da) of asparagine and glutamine amino acids, phosphorylation at serine, threonine, and tyrosine amino acids (+79.96633 Da) and N-terminus acetylation (+42.01057 Da) was set as variable modifications. To cover all 20 possible amino acid exchange cases at the **X** position, we performed searches with reference sequences covering all 20 standard amino acids as possible amino acid at the **X** position. We note that as leucine and isoleucine are isobaric, and we excluded isoleucine incorporation based on the amino acid identity of the utilized tRNAs. All cysteines were set to permanent no modification due to no alkylation procedure. An overall false discovery rate of 1% on both protein and peptide level was achieved by performing target-decoy database search using Percolator^92^.

### Quantification and statistical analysis

Statistical details regarding each experiment are described within the figure legends and in the *Results* section. For all mouse experiments, *n* refers to the number of mice in each experimental group. Animals were randomly assigned to treatment groups. Animal handling, fecal sample collection, DNA sequencing, and mass spectrometry raw data generation were blinded. All other forms of experiments and data generation were not blinded.

