## Supplementary_Material for "Firewalled synthetic commensal blocks horizontal gene transfer in the gut"

**Supplementary Figure 1.**

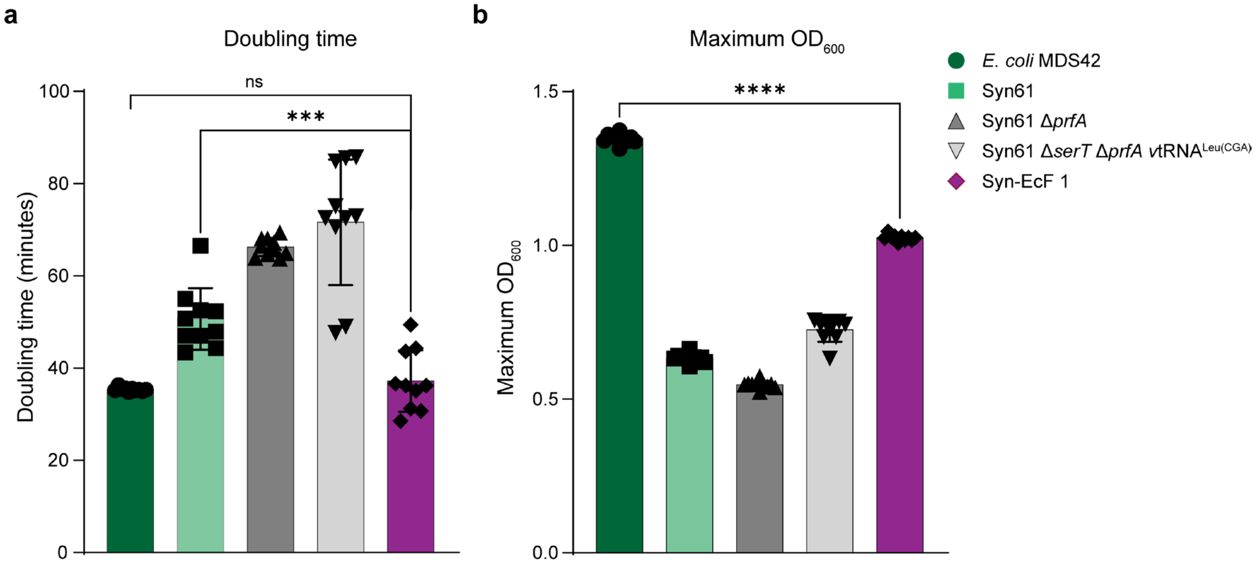


**Supplementary Figure 1. Doubling time (a) and maximal $OD_{600}$ (b) of the parental *E. coli* MDS42, Syn61, and strains created in this study under aerobic conditions**. We measured growth parameters aerobically in rich 2×YT broth at 37 °C. Data represent the mean; error bars denote SD based on *n*=10 independent biological replicates. *** indicates a P = 0.0006, **** indicates P ≤ 0.0001, while n.s. indicates a P = 0.468 based on two-tailed Mann-Whitney U test.

### Supplementary Table 1.

| Position | Mutation | Annotation | Gene | Functional annotation |
|---|---|---|---|---|
| **408501** | Δ2024 bp | | Δgsk–ybaL | gsk, ybaL deletion |
| **562075** | A→T | intergenic | gltI ← / ← lnt | glutamate/aspartate ABC transporter/apolipoprotein N-acyltransferase |
| **672775** | C→T | H3Y | bioB → | biotin synthase |
| **1022664** | C→T | G53E | phoP ← | DNA-binding transcriptional dual regulator PhoP |
| **1,293,766** | C→T | G216E | ydeP ← | putative oxidoreductase YdeP |
| **2,320,886** | Δ1,337 bp | | ΔygaZ | ygaZ deletion |
| **2,544,108** | T→A | T246L | prfB ← | peptide chain release factor RF2 |
| **2,876,758** | G→A | R314H | accC → | biotin carboxylase |
| **2,927,587** | A→C | L157* | rpsG ← | 30S ribosomal subunit protein S7 |
| **2,979,980** | ATCAGG ins | | nudE ← | ADP-sugar diphosphatase NudE |
| **3,143,455** | Δ1,004 bp | | ΔdppF–dppD | dppF, dppD deletion |
| **3,240,639** | T→C | L179P | kdtA → | 3-deoxy-D-manno-octulosonate transferase |
| **3,306,533** | T→A | Y76F | dgoR ← | putative DNA-binding transcriptional regulator DgoR |
| **3,555,503** | G→T | A264E | cytR ← | DNA-binding transcriptional repressor CytR |
| **3,590,695** | C→T | P99L | oxyR → | DNA-binding transcriptional dual regulator OxyR |
| **3,726,334** | Δ11 bp | intergenic | nrfG → / → gltP | nitrite reductase complex subunit NrfG/glutamate/aspartate: H(+) symporter GltP |

**Supplementary Table 1. Mutations observed in Ec-Syn$^{F}$ 1 following anaerobic gut-mimicking ALE.** Mutations were identified based on Illumina whole-genome sequencing followed by single-nucleotide polymorphism and indel identification using breseq (version 0.36.1)[3] compared to the parental genome. PrfB T246L mutation is induced by an ACG→TCG mutation, resulting in Leu in Syn-Ec$^{F}$ 1's genetic code.

**Supplementary Figure 2.**

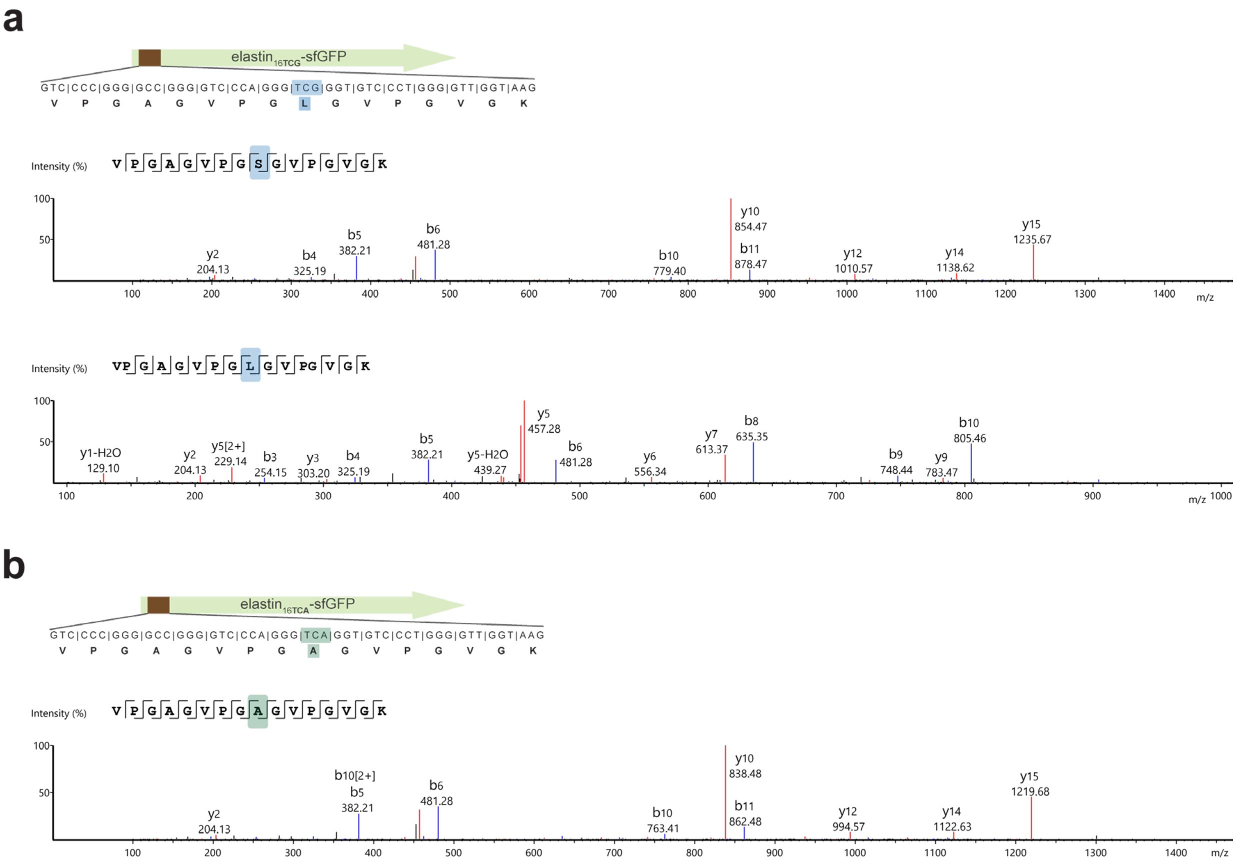


**Supplementary Figure 2. Genetic code of the evolved Syn61 Δ*serT* Δ*prfA* containing coliphage MF402939.1 tRNA$^{Leu(CGA)}$[1,2] insertion and the ALE-derived tRNA$^{Ala(UGA)}$, induced by an *alaT* anticodon TGC→TGA mutation**. The modified genetic code decodes (**a**) TCG codons as serine and leucine, while (**b**) TCA codons are translated as alanine. The amino acid identity of the translated TCG codon (**a**) and TCA codon (**b**) within elastin$^{(16TCA)}$–sfGFP–His$_6$ was confirmed by LC-MS/MS. The figure shows the amino acid sequence and MS/MS spectrum of the analyzed elastin$^{(16TCA/TCG)}$ peptide. LC-MS/MS data were collected once.

**Supplementary Figure 3.**

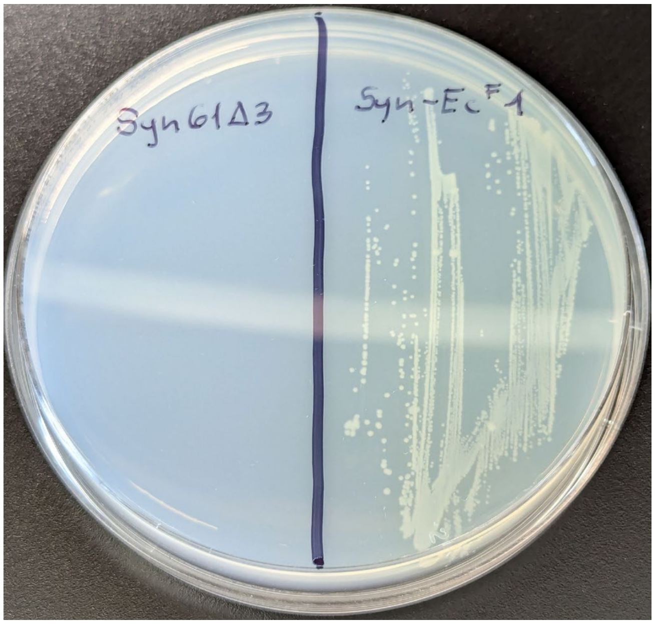


**Supplementary Figure 3. Growth of Syn-Ec$^{F}$ 1 on minimal M9 agar with glucose as the sole carbon source and the inability of Syn61Δ3 to form colonies on minimal M9 agar**. We streaked exponentially growing cultures of both strains side by side on M9 minimal salts agar plates with 1% glucose (Teknova M1200) and incubated them at 37 °C until colony formation.

**Supplementary Figure 4.**

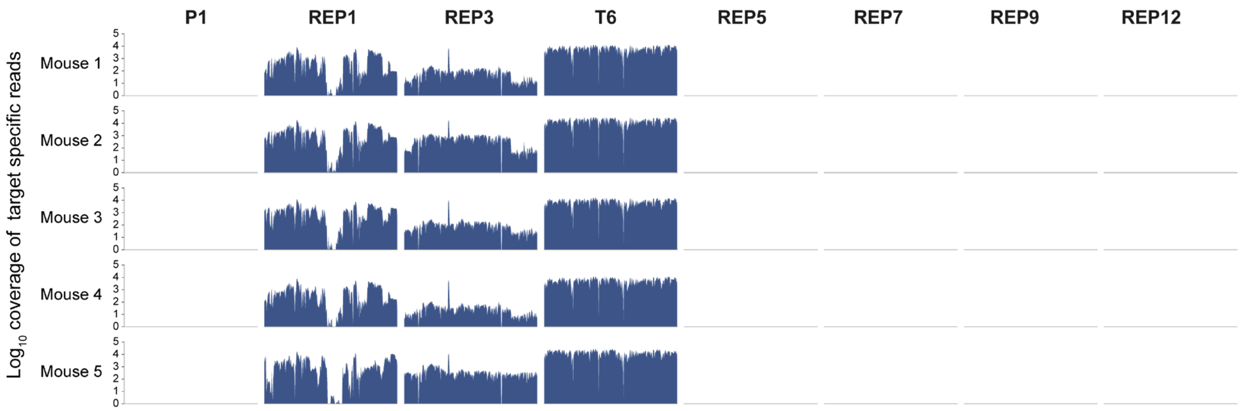


**Supplementary Figure 4. Next-generation sequencing read coverage of coliphage genomes in fecal metagenome samples of animals colonized with *E. coli* Nissle 1917**. The figure shows phage-unique Illumina NGS read coverage after 5 days of gut phage colonization in *E. coli* Nissle 1917-colonized mice. Phage-unique sequencing reads were identified as described in section "*Sequencing-based gut bacteriophage abundance analysis*" (Methods), allowing us to accurately distinguish the abundance of closely related bacteriophages with high genome-sequence homology.

**Supplementary Figure 5.**

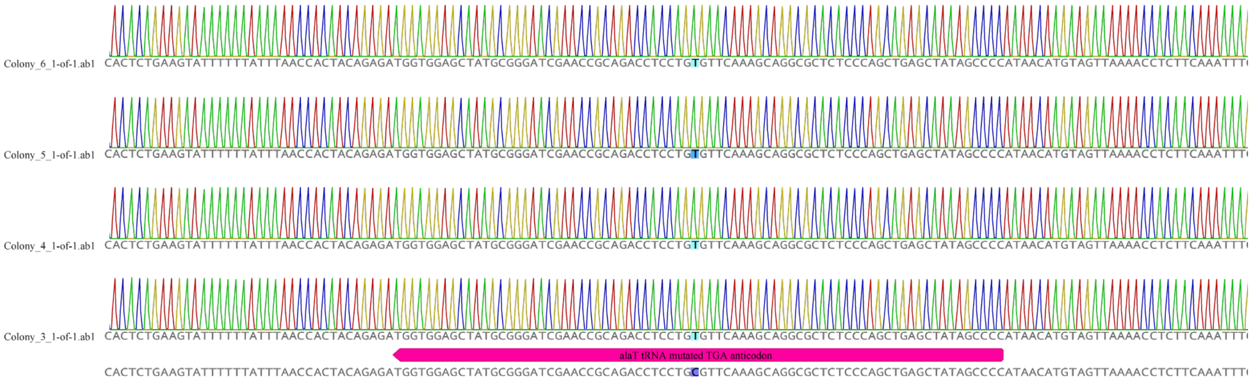


**Supplementary Figure 5. Nanopore sequencing of tRNA$^{Ala(UGA)}$ following experimental phage-cell co-evolution**. Four independently picked colonies were subjected to targeted Nanopore sequencing, and sequencing reads were aligned to the parental strain's genome. We note that performing the same analysis at the tRNA$^{Ala(CGA)}$ locus detected no mutations within tRNA$^{Ala(CGA)}$.

### Supplementary Table 2.

| Position | Mutation | Annotation | Gene | Functional annotation |
|---|---|---|---|---|
| 155466 | Δ54 bp | | dksA ← | RNA polymerase-binding transcription factor DksA |
| 2,420,342 | G→A | viral tRNA-Leu G5→A | | coliphage MF402939.1 tRNA-Leu(CGA) G5→A |
| 3,498,574 | C→T | M143I | yihP ← | 2,3-dihydroxypropane-1-sulfonate export protein |
| 3,501,145 | A→T | intergenic | yihQ ← / ← yihR | sulfoquinovosidase/putative aldose 1-epimerase YihR |
| 3,506,916 | Δ5 bp | coding (440-444/786 nt) | yihW → | DNA-binding transcriptional regulator YihW (CsqR) |

**Supplementary Table 2. Mutations observed in Syn-Ec$^{F}$ 1 following long-term within-gut evolution in mice.** Mutations were identified based on Illumina whole-genome sequencing followed by single-nucleotide polymorphism and indel identification using breseq (version 0.36.1)[3] compared to the parental genome.
